# Defects in autophagy lead to selective *in vivo* changes in turnover of cytosolic and organelle proteins in Arabidopsis

**DOI:** 10.1101/2021.04.29.441983

**Authors:** Lei Li, Chun Pong Lee, Akila Wijerathna-Yapa, Martyna Broda, Marisa S. Otegui, A. Harvey Millar

## Abstract

Identification of autophagic protein cargo in plants by their abundance in *autophagy related genes* (*ATG*) mutants is complicated by changes in both protein synthesis and protein degradation. To detect autophagic cargo, we measured protein degradation rate in shoots and roots of Arabidopsis *atg5* and *atg11* mutant plants. These data show that less than a quarter of proteins changing in abundance are probable cargo and revealed roles of ATG11 and ATG5 in degradation of specific cytosol, chloroplast and ER-resident proteins, and a specialized role for ATG11 in degradation of proteins from mitochondria and chloroplasts. Our data support a role for autophagy in degrading glycolytic enzymes and the chaperonin containing T-complex polypeptide-1 complex. Autophagy induction by Pi limitation changed metabolic profiles and the protein synthesis and degradation rates of *atg5* and *atg11* plants. A general decrease in the abundance of amino acids and increase in several secondary metabolites in autophagy mutants was consistent with altered catabolism and changes in energy conversion caused by reduced degradation rate of specific proteins. Combining measures of changes in protein abundance and degradation rates, we also identify ATG11 and ATG5 associated protein cargo of low Pi induced autophagy in chloroplasts and ER-resident proteins involved in secondary metabolism.

**Single Sentence Summary:** Protein cargo of autophagy in plants can be discovered by identifying proteins that increase in abundance and decrease in degradation rate in mutants deficient in autophagy machinery

## Introduction

Autophagy enables cellular sugar, lipid and protein recycling and maintenance through the trafficking of cellular material into the hydrolytic environment of the vacuolar lumen. Autophagic degradation involves general and selective processes and is controlled through both autophagy related genes (ATGs) and a range of receptor recognition mechanisms (An and Harper, 2018; Marshall and Vierstra, 2018). Large protein complexes like ribosomes, proteasomes, and protein aggregates are recognized through receptors-adaptor interaction and engulfed by autophagosomes for delivery to vacuoles (Marshall et al., 2015; Floyd et al., 2016; Jung et al., 2020). Autophagic degradation is also involved in clearance of chloroplasts, mitochondria, peroxisomes, and ER during developmental transitions or stress responses (Liu et al., 2012b; Farmer et al., 2013; Li et al., 2014; Khaminets et al., 2015; Izumi et al., 2017; Zhang et al., 2020).

ATG proteins participate in autophagosome induction, membrane delivery, vesicle nucleation, cargo recognition, and phagophore expansion and closure (Marshall and Vierstra, 2018). While some ATGs are encoded by single or duplicated genes in plants, there are notable exceptions like ATG8 and ATG18 which are encoded in multi-gene families (Thompson et al., 2005; Xiong et al., 2005; Yoshimoto et al., 2009a). In the multi-stage conjugation system that mediates phagophore formation, ATG8 and ATG12 are each typically conjugated to ATG7 and transferred separately to ATG3 and ATG10, respectively. Subsequently, ATG8 is covalently attached to phosphatidylethanolamine (PE) and ATG12 is attached to ATG5, forming an E3 ligase complex. The ATG5-ATG12 conjugate mainly contributes to phagophore expansion and maturation. *ATG5* mutants in Arabidopsis fail to form autophagosomes, show a general disruption in subsequent autophagy-related processes, and senesce under nitrogen- and carbon-limiting conditions (Thompson et al., 2005; Yoshimoto et al., 2009b). ATG11 is an accessory protein that aids the scaffolding of the ATG1 kinase regulatory complex to the expanding phagophore. ATG11 is reported to promote vesicle delivery to vacuoles by stabilizing the ATG1/13 complex, but does not appear to influence ATG12-ATG5 or ATG8-PE conjugates. Arabidopsis mutants deficient in *ATG11* also senesce rapidly under nitrogen- and carbon-limiting conditions and fail to degrade mitochondrial proteins during dark-induced senescence (Li et al., 2014; Li and Vierstra, 2014).

The apparent accumulation of specific sets of proteins in *atg* mutant plants (Avin-Wittenberg et al., 2015; McLoughlin et al., 2018; Have et al., 2019; McLoughlin et al., 2020) may be caused directly by a failure in autophagy-dependent protein degradation, or indirectly through an increase in protein synthesis rate due to their enhanced gene expression or translation. A failure in protein degradation could also be accompanied by lower levels of protein synthesis through either feedback attenuation of gene expression or translational control. Thus, using steady-state protein abundance as sole criterion to identify autophagic protein targets is prone to errors (Wijerathna-Yapa et al., 2021). A couple of multi-omics studies surveying the effect of autophagic recycling on proteome remodeling attempted to use comparisons of mRNA and protein abundance in maize *atg12* genotypes to resolve this issue (McLoughlin et al., 2018; McLoughlin et al., 2020). These studies found that more than half of the proteins that accumulated in *atg12* plants did not have consistent changes in the abundance of their mRNA. Using a similar approach, a lack of correlation in protein-transcript changes was also observed in *atg5* plants (Have et al., 2019). In addition, differential regulation of translational rates imply that the same amount of mRNA will not always result in the same level of translation, especially in autophagy mutants in which ribosome accumulation has been reported (Gretzmeier et al., 2017; McLoughlin et al., 2018). It therefore remains an open question as to which proteins that accumulate in *atg* mutants are actual autophagy cargo and which are autophagy-related changes in protein synthesis rate.

Autophagy deficiency also leads to changes in the abundance of metabolic intermediates in plants. Metabolic profiling shows that *ATG*-deficient mutants respond differently to prolonged darkness or nutrient limitation by undergoing extensive rearrangement in primary and secondary metabolism (Masclaux-Daubresse et al., 2014; Avin-Wittenberg et al., 2015; Barros et al., 2017; McLoughlin et al., 2018; Have et al., 2019; McLoughlin et al., 2020; Barros et al., 2021). These reports show that the changes in metabolic profiles and protein abundance in *atg* lines following nutrient limitation is complex, depends deeply on environmental, developmental, and tissue/organ context.

Our previous use of stable isotope progressive labelling to measure protein turnover rates in barley and Arabidopsis revealed that organelles and intra-organellar components are degraded at different rates (Nelson et al., 2014; Li et al., 2017). These rates resulted from the combined action of specific proteases in different subcellular compartments, the proteasome, and autophagy-dependent and autophagy-independent vacuolar degradation. In this study, we combined a quantitative analysis of changes in protein abundance with a stable-isotope progressive labelling strategy to measure protein degradation rates in *atg5* and *atg11* lines of Arabidopsis. Using these data, we quantified the contribution of autophagic degradation to the clearance of different organelles under control and Pi-limiting conditions. Changes in protein abundance and turnover rate provide clues to understand broad changes in metabolite levels in autophagy mutants, links between cellular trafficking and autophagic flux, and identity a range of autophagy target proteins in shoot and root tissues.

## Results

### Arabidopsis *atg5* and *atg11* mutants do not show accelerated senescence in hydroponics at early stages of leaf production

The early onset of leaf senescence after 35 days of growth, or following bolting, is a widely reported Arabidopsis phenotype in well-studied autophagy deficient mutants, including *atg5* and *atg11* (Yoshimoto et al., 2009b; Li et al., 2014). We choose an earlier developmental stage prior to leaf senescence for our experiments to avoid senescence-associated protein abundance and degradation rate changes from dominating our analysis. After visual inspection of hydroponically grown wild type (WT) and *atg* mutant plants we decided to use 21-day-old plants for analysis as no visible signs of senescence were observed in 10 rosette leaves which resembled growth stage 1.10 plants as reported previously (Boyes et al., 2001) (**Fig S1A)**. Consistent with the visible appearance of plants, the quantum efficiency of photosystem II (Fv/Fm) was the same in WT, *atg5,* and *atg11* leaves and no evidence of early senescence hot spots were observed in pulse-amplitude-modulation (PAM) fluorometry images (**Fig S1B-C**).

### Deficient autophagic flux leads to broad changes in the abundance of proteins in Arabidopsis roots and shoots

Changes in relative protein abundance between different genotypes and their biological replicates were measured using a ^15^N reference sample as a control. Total root or shoot proteins extracted from WT, *atg5* and *atg11* grown in ^14^N media were mixed with equal amounts of reference samples of ^15^N fully labeled WT shoot or root protein (Li et al., 2017). The combined samples were digested by trypsin and the resulting peptides fractionated and analysed by mass spectrometry. In total, 25,771 non-redundant peptides from root tissues and 18,939 peptides from shoot samples could be quantified using ratios of ^14^N sample peptides to ^15^N reference peptides. These peptides mapped to 1,265 non-redundant proteins in roots and 777 in shoots that could be quantitatively compared between WT and *atg* lines (**DataS1**). We performed pairwise comparisons between WT, *atg5* and *atg11* using protein sets that were quantified in all three biological replicates. Volcano plots showed that both autophagy mutants exhibited symmetric distributions for sets of proteins increasing or decreasing in abundance. Fold changes (FC) in protein abundance in *atg11*/WT show a relatively narrow range of FC (2-fold FC in root and 4-fold FC in shoot) compared with a wide range of FC in *atg5*/ WT protein abundance (4-fold FC in root and 8-fold FC in shoot) **(FigS2).**

To dissect the role of autophagy in protein homeostasis in different cellular compartments, we displayed the distributions of relative change in protein abundance according to the known subcellular localization of each protein (Hooper et al., 2014; Hooper et al., 2017) (**Fig1, DataS2**). Proteins located in the cytosol and peroxisomes of both shoots and roots showed higher median abundance in *atg11* and *atg5* mutants than in WT. Conversely, proteins in the nucleus, plasma membrane, vacuoles, and those secreted to the extracellular space, showed lower median abundances in the autophagy mutants compared to WT. The majority of the 53 plastid proteins found in roots showed lower abundance in the mutants, but an overall increase of chloroplast protein abundance was observed in shoot tissue from both mutant lines (**Fig 1**). ER-resident proteins were more abundant in the root but not in shoot of the mutants. A higher abundance of mitochondrial proteins was found in both shoots and roots of *atg11,* but only in the shoot of *atg5* compared to WT.

**Fig 1.**
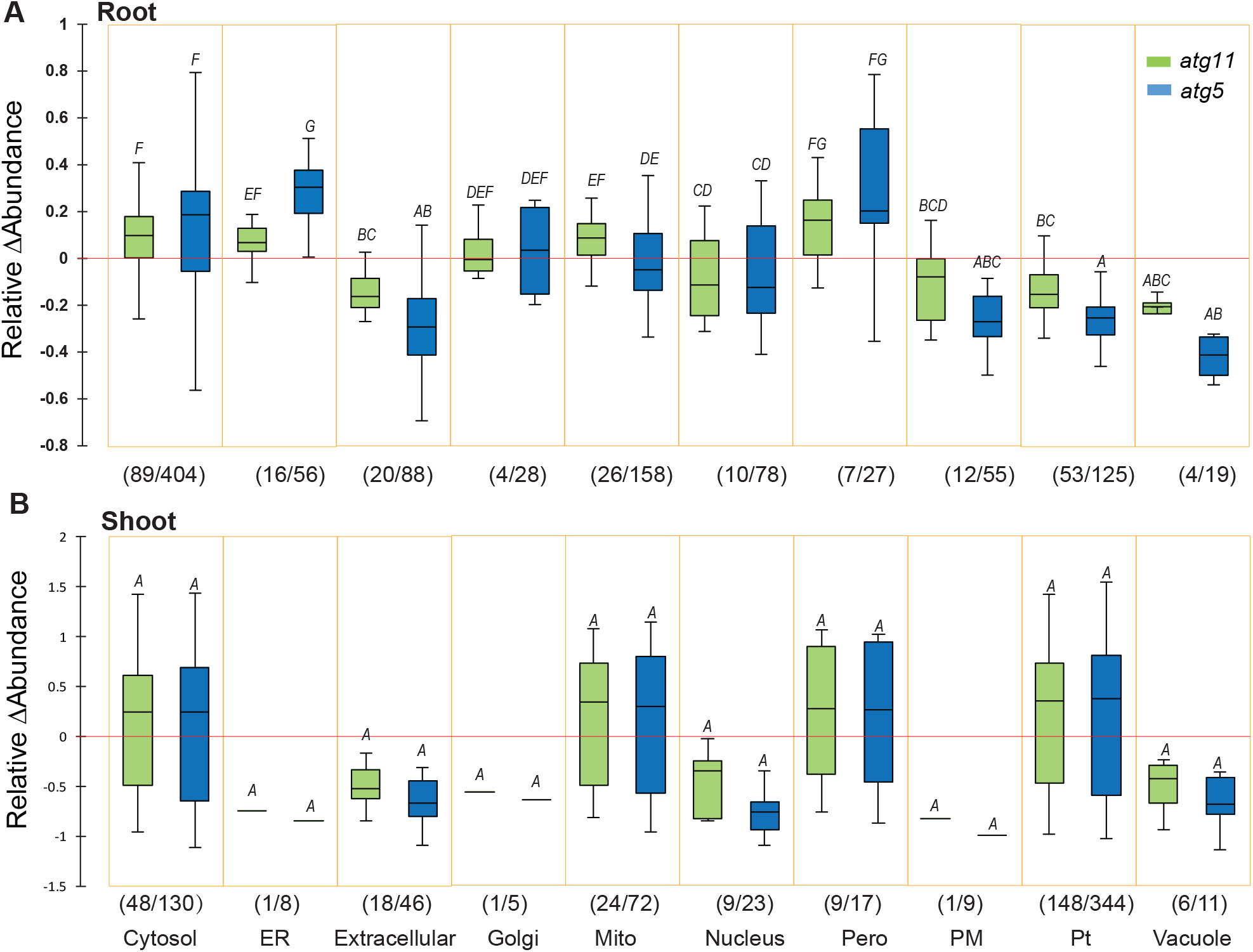
Changes in abundance of proteins in *atg5* and *atg11* that are resident in different subcellular locations. Box plots of relative changes in abundance of 241 root (**A**) and 265 shoot (**B**) proteins from *atg5* and *atg11* which significantly changed in abundance in comparison with WT (p<0.05). These were from the larger set of 1114 root and 698 shoot proteins that were quantified in all genotypes. ΔAbundance changes of specific root and shoot proteins are shown in **Fig S2** and **Fig S3**. Numbers of significantly changing (x) and total quantified (y) proteins for each subcellular location (x/y) are shown. A comparison of k-samples distributions (Kruskal-Wallis) was performed in XLSTAT to evaluate the level of changes in subcellular locations. Changes in abundance of proteins in root can be divided into A-G groups with increasing values. Changes in abundance of proteins in shoot can be only divided into A group. An explanation of values defining each group would be important.

To further understand the role of autophagy in regulating the protein synthesis and degradation machineries, we investigated abundance changes in ribosomal and proteasomal subunits in both mutants compared to WT (**FigS3, DataS2**). For ribosomes, 70-80% of r-subunits showed a trend of higher abundance in the mutants, with 17 out of 61 r-protein in root and 2 out of 7 ribosomal r-proteins in shoot showing statistically significant increases (Student’s T-test, P<0.05). More than half of the proteasomal subunit proteins identified also tended to be more abundant in both mutants, with 2 out of 23 proteasomal subunit proteins in root and 1 out of 6 in shoot showing statistically significant increases (Student’s T-test, P<0.05).

### Specific proteins changed in abundance in a manner unexpected for their function or subcellular location in autophagy-deficient plants

To investigate specific protein abundance changes in root (**Fig S4**) and shoot (**Fig S5**), proteins with statistically significant changes in *atg5* and *atg11* were then categorized by their subcellular localizations and functions.

In roots, the cytosolic Chaperonin Containing T-complex polypeptide-1 (CCT) protein complex subunits, ribosomal subunits, enzymes of amino acid metabolism (GAD1, ASP2, MMT and OLD3) and glycolytic enzymes accumulated in *atg11* and *atg5* **(Fig S4)**. In contrast, cytoskeleton-related proteins including villins (VLN4), actin (ACT7) and tubulin (TUB2,4,6,8,9), enzymes of amino acid metabolism (MAT3 and BCAT4), and phosphatidylinositol transfer proteins (At1g30690 and At1g72160) from the secretory pathway showed decreased abundance in both *atg11* and *atg5*. Eleven mitochondrial proteins, including components of the electron transport chain and TCA cycle, showed increases in abundance, several mitochondrial stress response proteins, such as mtHsc70-1, mtHsc70-2 and GPX6, displayed a decreased abundance in both mutants. Ten mitochondrial proteins (including ATP synthase beta subunit, CPN10, ATPHB3, TOM5 and carbonic anhydrase) showed different patterns in *atg11* and *atg5*, with their abundance typically increased in *atg11* but decreased in *atg5* **(Fig S4).** We also found that in roots, proteins involved in vesicle transport specifically accumulated in *atg5* but not in *atg11*; these proteins included clathrin heavy chain1 (At3g11130) associated with plasma membrane and Golgi, and the coatomer alpha, delta and gamma-subunits (At2g21390, At5g05010 and At4g34450) of the COP1 coat, which is required for intra Golgi-transport, retrograde transport from Golgi to ER, and Golgi maintenance. ER-resident proteins, such as AtBAG7(At5g62390), CNX1 (At5g61790) and PDIL1-3(At3g54960), also show a higher abundance in *atg5* than *atg11* when compared to WT (31% in *atg5* vs 6% in *atg11*).

Different sets of proteins were quantified in shoots compared with roots due to the variation in their absolute abundance in photosynthetic and non-photosynthetic tissues. In shoots, almost half of quantified cytosolic proteins with significant changes in abundances were less abundant in mutant lines **(Fig S5)**. Similar to the protein set from the roots, cytosolic ribosomal subunits, enzymes of amino acid metabolism (MAT3 and ATCIMS) and glycolytic enzymes from shoots showed increased abundance, while profilin1 and profilin2, which regulate the organization of actin cytoskeleton, showed reduced abundance in both *atg11* and *atg5*. Peptidylprolyl isomerase enzymes (FKBP12, ROC1, ROC3 and ROC5) and proteins with redox activity (TRX3, TPX1 and CSD1) showed decreased abundance in both *atg11* and *atg5* shoots. In shoots, the mitochondrial redox proteins (GPX6, PRXIIF), CPN10, and membrane-localized electron transport chain subunits showed decreased abundance while TCA cycle enzymes and matrix-localized ETC subunits accumulated in both *atg11* and *atg5*.

In chloroplasts, most stromal proteins showed increased abundance, while photosystem II (PSII) subunits, photosystem I (PSI) reaction center (PSAN), cytochrome *b_6_/f* (PetC), plastocyanin (DRT112 and PETE1), thioredoxins, and protein folding associated proteins were less abundant in both *atg11* and *atg5* **(Fig S5)**. Most quantified shoot plastid proteins showed consistent changes in abundance in both mutant lines with few exceptions.

To determine whether these many changes in the abundance of specific root proteins were reflected in changes in the cellular architecture, we analysed the ultrastructure of root tips of 24-day-old WT, *atg5*, and *atg11* plants processed by high-pressure freezing/ freeze-substitution and resin-embedding. In longitudinal sections of root tips, we identified two areas of interest, the meristematic area (up to 100 microns from the quiescent center towards the elongation zone) and the area where cells started to develop large vacuoles (between 100 and 200 microns from the quiescent center) (**Fig 2 A-C**). We imaged multiple middle sections of two roots of each genotype and measured the cell area and the area occupied by the nucleus, mitochondria, and vacuoles as well as the tonoplast length per section (**Fig 2 D-H**). We did not find statistically significant differences in any of these parameters between WT and *atg* mutants; however, there were consistent trends showing slight increase in vacuole surface and a reduction in tonoplast length/perimeter in the two *atg* mutants, in actively vacuolating cells. These results indicating that the changes in the proteome of *atg5* and *atg11*, did not induce drastic changes in the cellular organization of mutant root cells.

**Fig 2.**
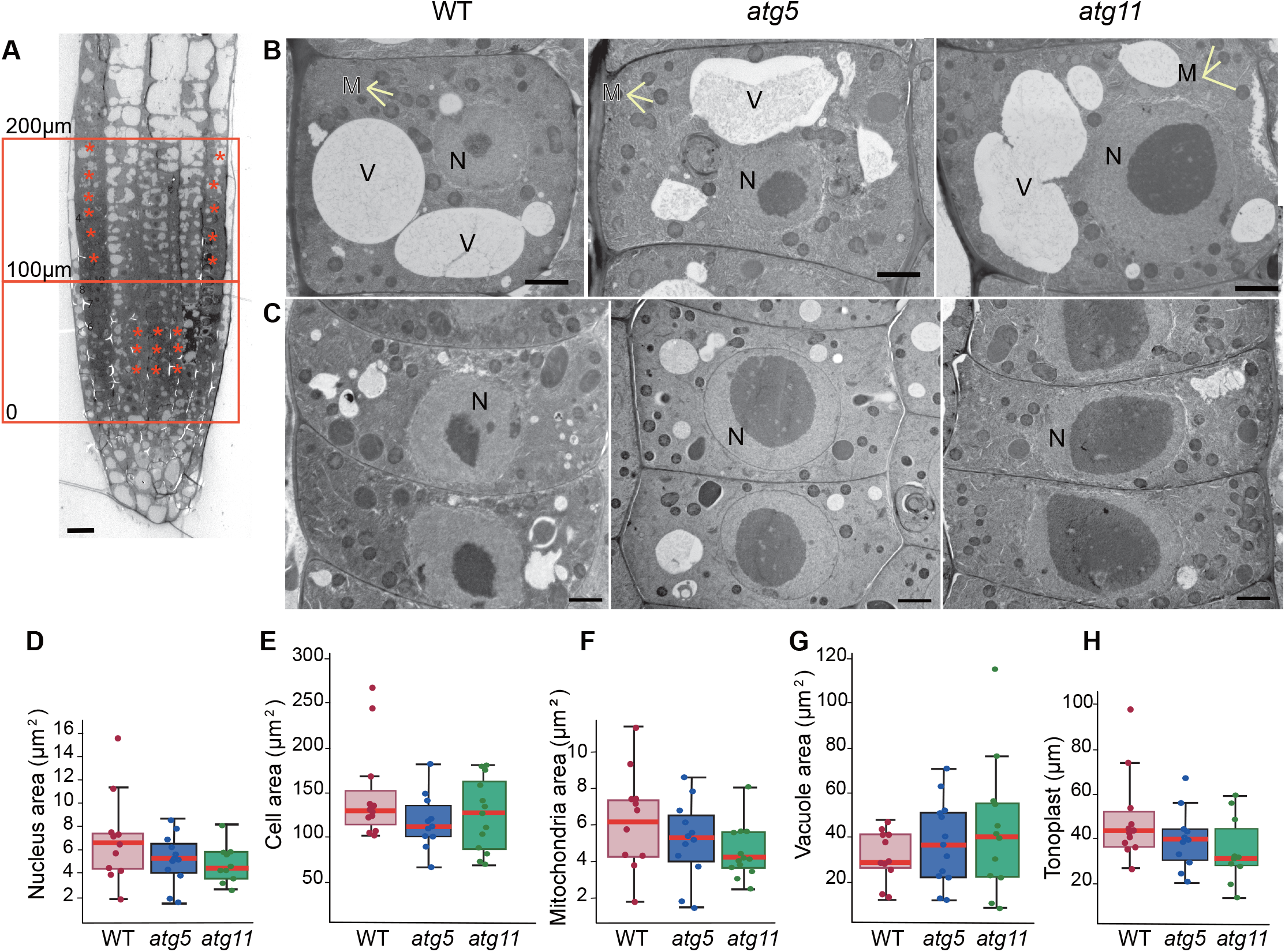
Transmission electron microscopy analysis of WT, *atg5*, and *atg11* root cells. (A) Longitudinal section of a WT root showing the areas selected for analysis: meristem region (up to 100 μm from quiescent center; QC) and the adjacent area up to 200 μm from the QC where cells are actively developing vacuoles. Asterisks indicate examples of cells that were analyzed. (B) Cell with developing vacuoles (C) Meristematic cells; D-E Quantification of vacuolated cell area per section (D), nuclear area in meristematic cells (E), mitochondrial and vacuolar area per section of vacuolated cells (F,G) and length of tonoplast per section of vacuolated cells (H). Between 10 and 13 cells from two roots of each genotype were used for this analysis. M, mitochondria,; N, nucleus; V, vacuole. Scales bars= 10 μm in (A); 2 μm in (B,C).

However, we noticed that approximately 25% of the cells in the *atg5* root tips contained abnormal Trans-Golgi networks (TGN) with largely dilated bulges or vesicles (**Fig 3 A-B**) and large concentric membranous systems (**Fig 3 C-F**). In some cases, the edges of these abnormal large membranes had bulges and budding profiles reminiscent of Golgi/TGN cisternae (**Fig 3C and D**). In some other examples, we were able to image coats assembled on budding sites on the membrane edges (**Fig 3E**, arrowheads), which is consistent with the abnormal accumulation of COP1 coatomer subunits and clathrin in *atg5*. Whereas most of these structures seem to enclose ribosomes and cytoplasm, approximately 10% of them displayed rounded electron dense aggregates 2-3 times larger than a ribosome (**Fig 3E**).

**Fig 3.**
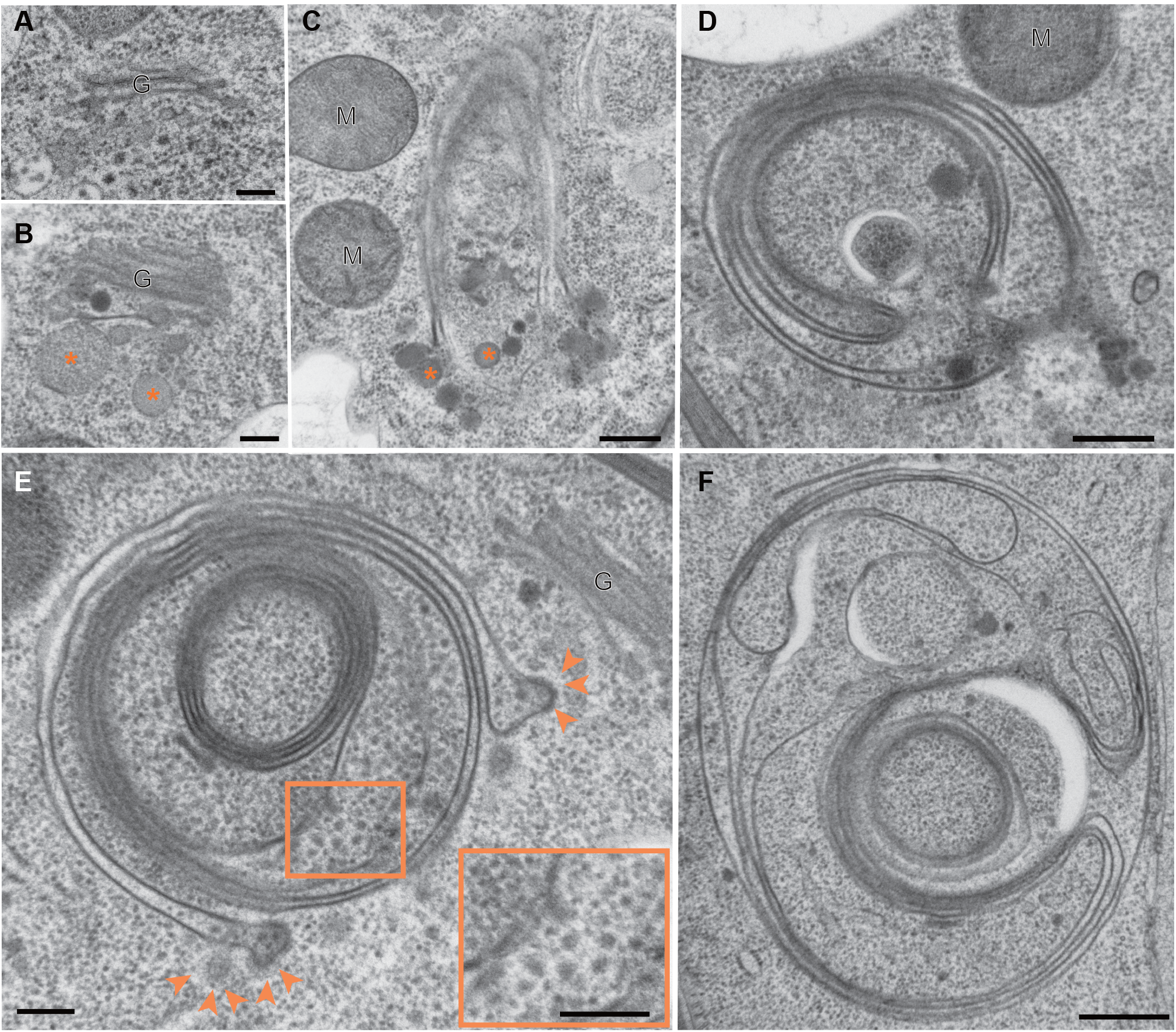
Abnormal organelles in *atg5* root cells. (A) Golgi stack in WT cells. (B) Golgi stack and associated TGN with dilated vesicle profiles (asterisks) in *atg5.* (C)-(F) Large membranous structures with concentric membranes in *atg5*. Some of these structures displayed budding profiles at their edges (asterisks in C). Budding sites with assembled coats (E, asterisks) are commonly seen on these structures. Enclosed by these membranes, there are electron dense aggregates 2-3 times larger than a cytosolic ribosomes (E and inset). G, Golgi stack; M, mitochondrion. Scale bars= 200 nm (A,B, E); 400 nm (C,D); 500 nm (F).

### Autophagy deficiency changes the degradation rate of specific organelle proteins in Arabidopsis root and shoot

To compare specific protein abundance changes with changes of specific protein degradation rates, we utilized a ^15^N progressive labelling strategy (Li et al., 2017) to quantify protein degradation rates in the three genotypes. For this, the media of hydroponically grown plants was switched from ^14^N to ^15^N nutrient salts to label newly synthesized proteins over three days, and the fraction of each peptide that was ^15^N-labelled (Labelled Peptide Fraction; LPF) was calculated using peptide mass spectrometry. In total, LPF for 11,179 peptides in roots and 7,145 peptides in shoots was quantified in three biological replicates across the three genotypes. From these LPFs, the degradation rates (K_D,_ d^−1^) of 558 root proteins and 505 shoot proteins were obtained (**Data S3**) and relative changes in K_D_ values were visualize by volcano plots (**Fig S6**). In roots, most proteins with slower degradation rates in *atg11* (68%) and *atg5* (82%) were located in the cytosol, followed by smaller proportions that were located in mitochondria and ER **(Table 1)**. In shoots, proteins that degraded slowly were predominantly from the cytosol, chloroplasts, and mitochondria. A higher proportion of mitochondrial proteins with slower degradation rates were detected in *atg11* roots and shoots (17%, 21%) compared to *atg5* (2%, 0%). There was also a higher proportion of chloroplastic proteins with slower degradation rates in shoots of *atg11* (33%) compared to shoots of *atg5* (7%).

**Table 1.** Cellular compartments with significant slower protein degradation rate in *atg5/11* mutants compared with wild-type Arabidopsis. Percentage of proteins resident in major cellular compartments that showed significant slower degradation rate in *atg11* and *atg5* than in Wt. In root, the majority of proteins with slower degradation rate are located in the cytosol, mitochondrion or ER. In shoot, the majority are in the cytosol, chloroplast and mitochondrion. Cellular localization of proteins beyond the top three are marked as others. Numbers in brackets are the number of proteins with slower protein degradation rate divided by the total number of proteins from that location that were analysed.

|  | *atg11/Wt* | *atg5/Wt* | Subcellular location |
| --- | --- | --- | --- |
| **Root** | 68% (45/66) | 82% (42/51) | Cytosol |
|  | 17% (11/66) | 2% (1/51) | mitochondrion |
|  | 5% (3/66) | 4% (2/51) | ER |
|  | 11% (7/66) | 12% (6/51) | Others |
| **Shoot** | 33% (8/24) | 64% (9/14) | Cytosol |
|  | 33% (8/24) | 7% (1/14) | chloroplast |
|  | 21% (5/24) | 0% (0/14) | mitochondrion |
|  | 13% (3/24) | 29% (3/14) | Others |

In roots, proteins with significant slower degradation rate in both *atg5* and *atg11* **(Fig 4)** included forty cytosolic proteins, two ER proteins (the chaperones CRT1 and CNX1), and one mitochondrial protein (ATP synthase D chain). Cytosolic proteins in this list can be broadly placed into three major functional categories: metabolism, ribosome subunits, and glycolytic enzymes. In shoots, twelve cytosolic proteins showed slower degradation rates in both *atg5* and *atg11*. We also found proteins with slower rates of protein degradation but with statistical significance only in one of the two mutants (**Table S1**). One example from this group was RPN10, which has been reported to be an autophagy receptor for the proteasome (Marshall et al., 2015). In contrast, four mitochondrial and eight chloroplastic proteins showed very different changes in degradation rate between *atg5* and *atg11*. These proteins show slower degradation rate in *atg11*, but no change or faster rates of degradation in *atg5*. These patterns suggested a specialized role of ATG11 in mitochondrial and chloroplast protein homeostasis.

**Fig 4.**
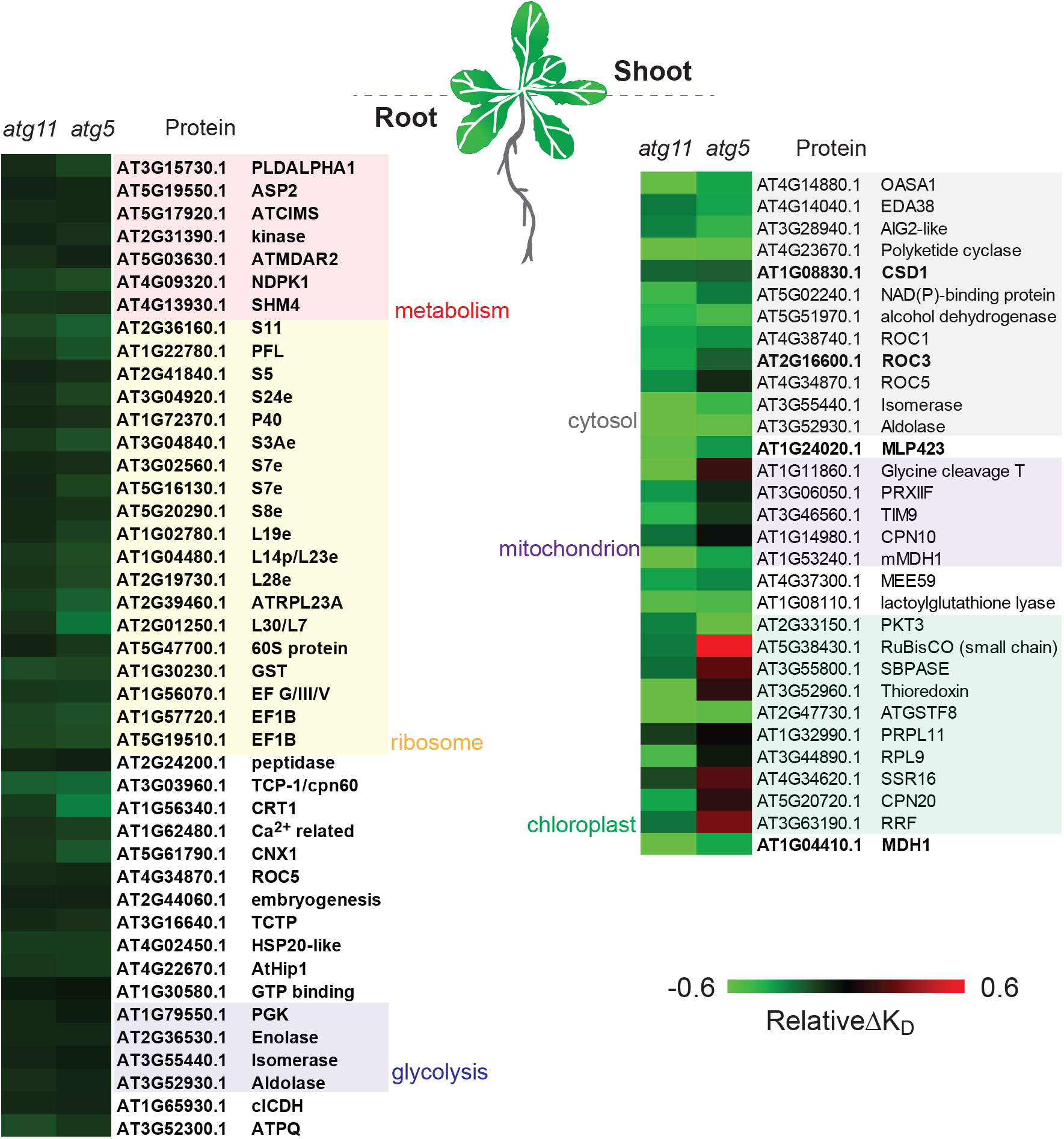
Specific proteins that degrade more slowly in *atg5 and atg11* roots and shoots compared with wild-type Arabidopsis. **(A)** A heatmap of 43 root proteins with significant slower degradation rate (relative ΔK_D_) in both *atg5* and *atg11*. (B) A heatmap of 31 shoot proteins with significant slower ΔK_D_ in *atg5* or *atg11*. Proteins with significance differences in both *atg5* and *atg11* and shown in bold font. Proteins are group according to the top three functional categories in root and top 3 organelles in shoot tissues. Specific protein degradation rates in WT, *atg5* and *atg11* can be found in Data S3.

### Identification of ATG5 and ATG11 targets from the combined protein degradation rate and abundance data

Protein abundance and degradation rate changes were then plotted orthogonally to pinpoint probable autophagy protein targets **(Fig 5)**. When *atg11* and *atg5* were compared to WT, we found that 140 and 200 root proteins and 116 and 187 shoot proteins showed significant changes in abundance and/or degradation rate (Student’s T-test, P<0.05). The response of these proteins could be grouped into the four quadrants with different responses as explained in **Fig 5**.

**Fig 5.**
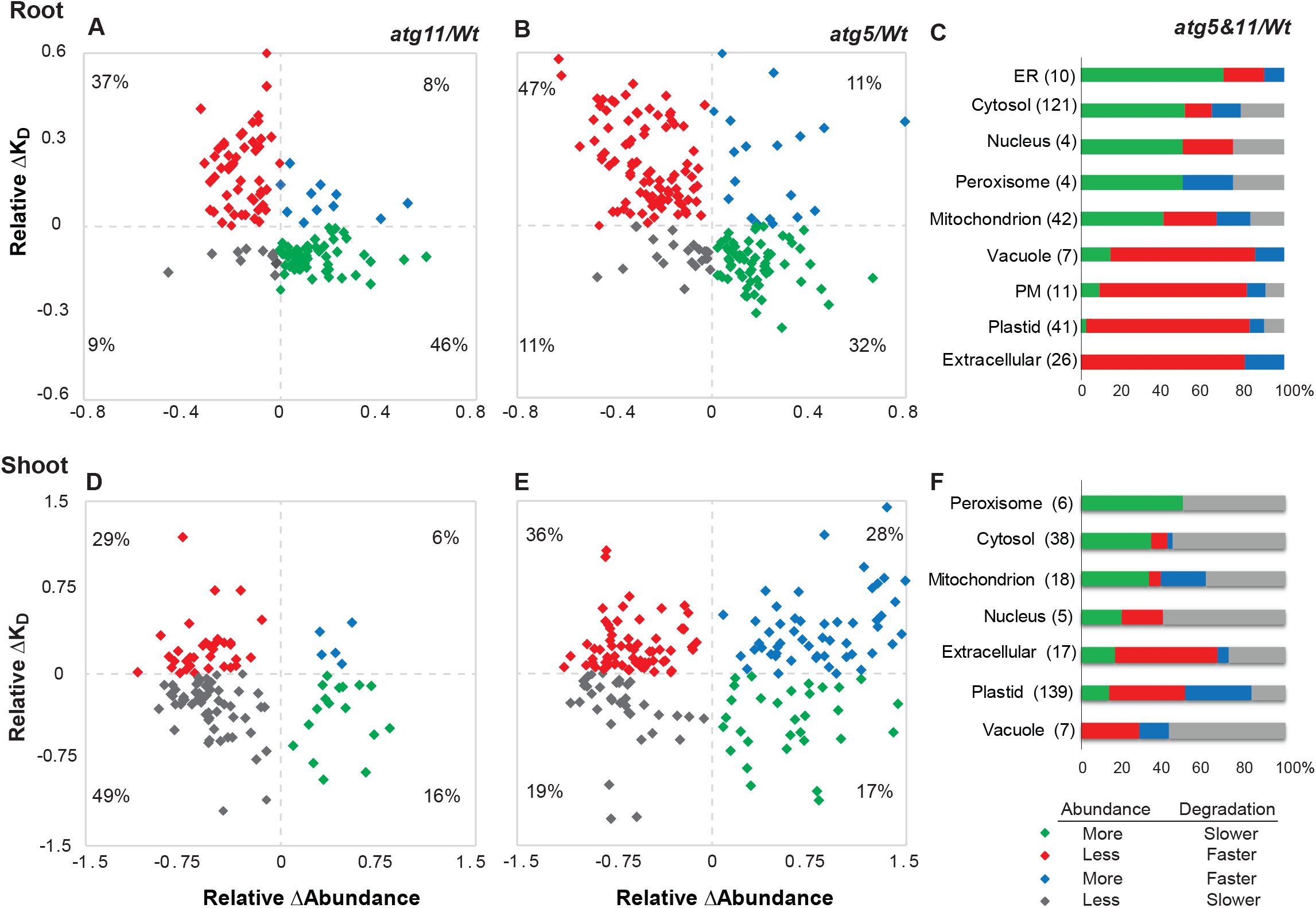
Combination of changes in abundance and degradation rate for proteins in *atg5 and atg11* from different cellular compartments. Matching sets of protein degradation rate changes (Relative ΔKD) and protein abundance changes (Relative ΔAbundance) were graphed orthogonally to identify putative autophagy cargo. 140 root proteins and 116 shoot proteins with significant changes in abundance or degradation (Student’s T-test, P<0.05) in *atg11* are plotted in **A** and **D**. 200 root proteins and 187 shoot proteins with significant changes in abundance/degradation (Student’s T-test, P<0.05) in *atg5* are plotted in **B** and **E**. The proportion of proteins shown in each quadrant that reside in a particular subcellular location in either mutant is shown for root data in **C** and shoot data in **F**. The four colors represent the four quadrants. Group 1 (green) represents proteins with slower degradation rates and greater abundance, consistent with direct changes driven by deficient autophagy substrate degradation; Group 2 (red) represents proteins with faster degradation rates and a lower steady-state abundance, potentially driven by alternative degradation pathways compensating for defects in autophagy; Group 3 (blue) contains proteins with faster degradation rates and greater abundance (likely driven by enhanced protein synthesis); and Group 4 (gray) contains proteins with slower degradation rates and lower abundance, putative examples of feedback regulated response to impaired autophagy degradation triggering decreasing protein synthesis.

Of the significantly changed proteins in roots, 80% were more abundant and slow degrading (Group 1-green) or less abundant and faster degrading (Group 2-red) **(Fig 5 A,B)**. The former are potential autophagy targets, and constituted more than half of the significantly changed proteins in *atg11* but less than 20% in *atg5* (**Data S5**). These proteins are typically localized to the ER, cytosol, nucleus, peroxisomes, and mitochondria (**Fig 5C**). Furthermore, they are components of mitochondrial oxidative phosphorylation, amino acid metabolism, glycolysis, the ribosome and proteasome, TCP-1 chaperones, and protein folding and processing in the ER **(Table S1)**. In comparison, most of the proteins that were degraded faster but accumulated less in the mutants (Group 2-red) were potential components of alternative and/or compensatory pathways and were localized to vacuoles, plasma membrane, plastids, and apoplast. From these, it was apparent that the mitochondrial TCA cycle and oxidative phosphorylation proteins showed the most distinct differences between the mutants, with 17 mitochondrial proteins belonging to Group 1 in *atg11* but not in *atg5*.

In shoots, only 50% of proteins that significantly changed were in Group 1 and 2, and Group 1 accounted for less than 20% of proteins in both mutants **(Fig 5D, E)**. In *atg11,* most of the remaining proteins were in Group 4 while in *atg5,* a higher proportion of proteins were in Group 3. Group 1 from shoots included mainly resident peroxisome, cytosolic, mitochondrial and chloroplastic proteins, with a smaller proportion of nuclear and vacuolar proteins **(Fig 5F)**. Proteins in the chloroplast showed different responses between mutant lines; ten chloroplastic proteins, including RUBISCO large subunit, fell into Group 1 in *atg11* but in Group 3 in *atg5* **(Table S2).** This again was consistent with ATG11 playing a specialized role in mitochondrial and chloroplastic protein degradation. A high proportion of proteins from shoots in Group 2 localize to plastids, vacuoles, or the apoplast **(Fig 5D,E)**. The higher proportion of proteins that fell into Group 4 in shoots compared to roots (**Fig 5C,F**) suggests that protein synthesis attenuation might mask autophagy degradation in shoot tissues.

### Pi limitation induces autophagy and changes metabolite abundances in hydroponically grown Arabidopsis

Nitrogen, phosphate or carbon limitation are reported to activate autophagy, promote cellular content degradation in plants and lead to early senescence in autophagy mutants (Yoshimoto et al., 2009b; Avin-Wittenberg et al., 2015; Barros et al., 2017; McLoughlin et al., 2018; Naumann et al., 2019; McLoughlin et al., 2020). However, it is unclear if such conditions lead to a generic induction of autophagy or of selective autophagy of stress-related targets. Nitrogen limitation conditions would limit our ability to use a ^15^N labelling and darkness would limit both carbon and ^15^N incorporation into amino acids (Nelson et al., 2014). Therefore, we subjected plants to Pi starvation to investigate its effect on protein abundance and degradation rate in all three genotypes **(Fig 6)**.

**Fig 6.**
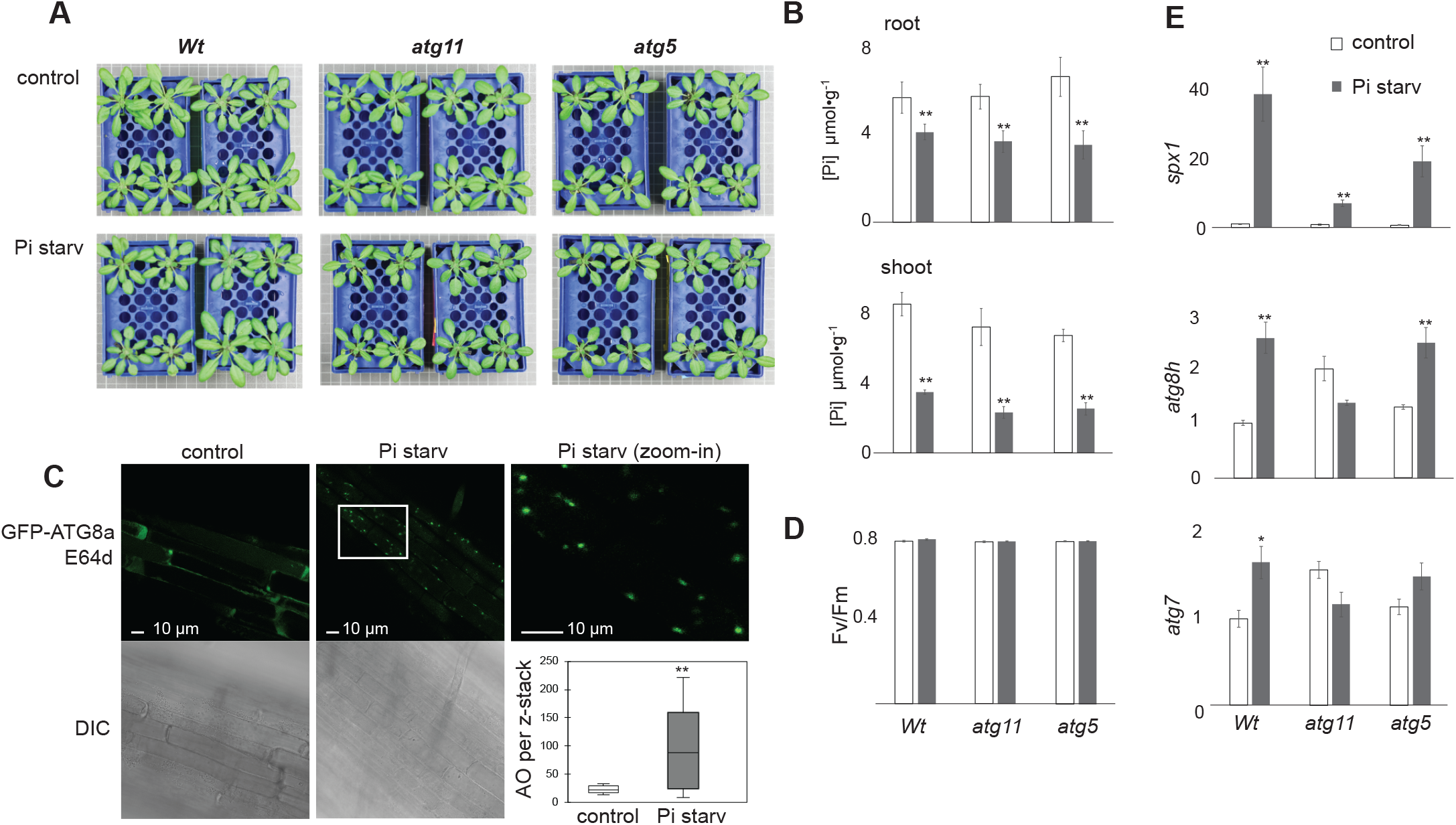
Pi limitation induces changes in *atg5*, *agt11* and WT Arabidopsis plants. (**A**) Arabidopsis plants grown in hydroponics for 21 days were transferred to fresh growth media with/without Pi for three days. (B) Free inorganic Pi concentration per fresh weight of root and shoot tissues was measured by a colorimetric assay method. (C) Root tips of a *GFP-ATG8a* line under control and Pi starvation conditions were treated with E64d (protease inhibitor) overnight before confocal imaging. Confocal image of elongation region are presented as well as differential interference contrast (DIC) images. Number of autophagic organelles (AO) per z-stack in roots grown under control or Pi limitation conditions (Kolmogorov-Smirnov two distribution test, **P<0.01). (D) Shoot tissue quantum efficiency of photosystem II (Fv/Fm) in *Wt*, *atg5 and atg11* lines. (E) Transcript abundance of *SPX1, ATG8H* and *ATG7* in *Wt*, *atg5 and atg11* under both control and Pi starvation conditions. Student’s T-test, *P<0.05, **P<0.01. Error bars show standard deviations of four biological replicates.

No visible phenotypic changes were observed in plants grown under Pi-limited conditions over 3 days of treatment (**Fig 6A)**, although both root and shoot Pi content was significantly reduced in all genotypes (**Fig 6B**). To monitor autophagy induction and autophagic flux, we performed an imaging analysis of a line expressing *GFP-ATG8a,* which is localized to autophagic membranes and autophagosomes in root cells. Abundant GFP-ATG8a-decorated organelles were evident in the elongation zone of Pi-limited roots but not in the equivalent root zone from control plants (**Fig 6C**). In shoots, the Fv/Fm ratio remained at 0.8 in all three genotypes under control and Pi-limiting conditions (**Fig 6D**). Consistent with the reduced Pi content, the transcript of the Pi sensor *SPX1* was induced in all three genotypes when plants were grown under Pi-limiting conditions (**Fig 6E**). Autophagy associated genes, *ATG8H* and *ATG7,* were also induced under limited Pi in WT and *atg5* but not in *atg11* plants (**Fig 6E**). Extension of Pi-limited conditions to >10 days led to purple coloration of rosette leaves, indicating stress-induced anthocyanin accumulation.

To further assess the impact of Pi limitation on metabolism, we performed a mass spectrometry-based profiling of selected primary and secondary metabolites in shoots and roots both between WT and autophagy mutants and within each genotype (**Fig 7, Data S11**).

**Fig 7.**
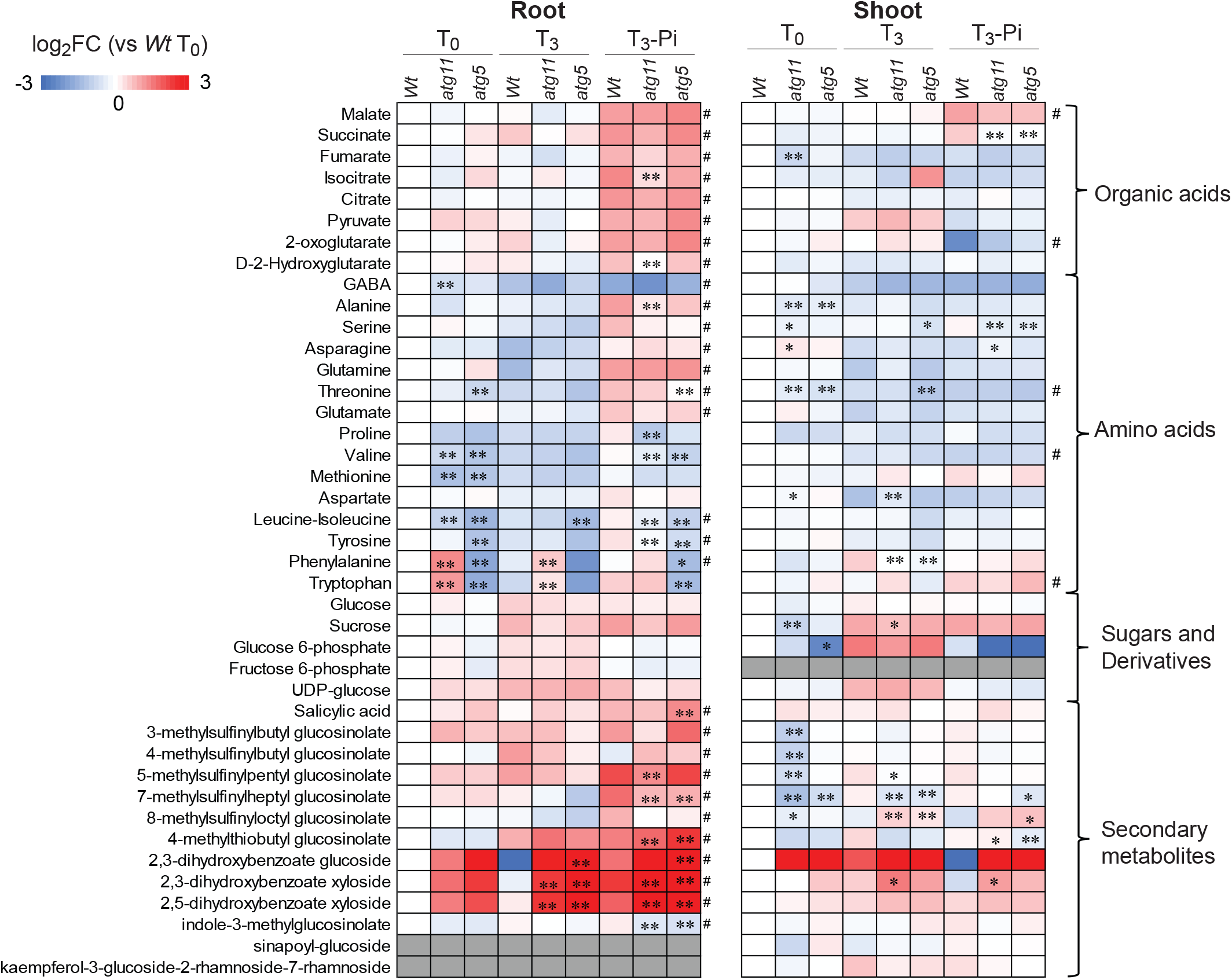
Primary and secondary metabolite profile changes under control conditions and Pi limitation in *atg5* and *atg11*. Heatmaps of changes in metabolite abundance in shoots and roots are shown. Metabolites were determined by LC/MS. Two-way ANOVA analyses were carried out to determine the genotype and Pi starvation effects on metabolite abundance changes. Metabolite content significantly altered due to Pi starvation, regardless of genotype, was labelled by a hashtag (#) (p< 0.05 for both T0 vs T3-P and T3-P vs T3+P). Metabolite content significantly altered due to genotype was determined by Tukey Ad hoc analysis for Col-0 vs either *atg5* or *atg11* and labelled by an asterisk (*); *p< 0.05 or ** p < 0.01. Metabolites with unquantifiable abundance in a given sample are shown in grey.

In roots, there was a drastic decrease of amino acids commonly reported to be highly responsive to nutrient limitation conditions (Araújo et al., 2011; Masclaux-Daubresse et al., 2014; Avin-Wittenberg et al., 2015; Barros et al., 2017). However, we observed very little effect on the abundance of amino acids and sugars in both mutants upon Pi limitation. Organic acids in roots generally did not change in abundance except for isocitrate and D-2-hydroxyglutarate, which were slightly more abundant in *atg11* under Pi starvation. Several glucosinolates changed in abundance in roots of autophagy mutants only under Pi limitation. Salicylic acid (SA) levels did not change in mutants under control conditions but accumulated in *atg5* under limited Pi (Fig 7), as also previously reported in dark-induced senescence (Yoshimoto et al., 2009b). Interestingly, SA-sugar conjugates, including 2,3-dihydroxybenzoate glucoside 2,3-dihydroxybenzoate xyloside, and 2,5-dihydroxybenzoate xyloside accumulated in autophagy mutants under both control and Pi-limiting conditions. SA conjugation inactivates SA; the accumulation of these compounds in autophagy mutants might be part of a mechanism to partially prevent the SA-dependent early senescence typical of autophagy mutants (Yoshimoto et al., 2009b).

In shoots, the levels of amino acids alanine, threonine, serine, and phenylalanine were reduced in the mutant lines. Asparagine and aspartic acid showed some accumulation in *atg11* but not in *atg5* under both control and Pi-limiting conditions. Organic acids did not show significant changes in abundance, with the exception of fumarate, which was reduced in *atg11*, and succinate, which decreased in both autophagy mutants but only under Pi limitation. Few changes in sugar and sugar derivative abundances were present in shoots, with glucose and glucose-6-phosphate decreasing slightly in *atg11* or *atg5*, respectively. All glucosinolates, except 8-methylsulfinyloctyl glucosinolate, decreased in abundance in shoots of autophagy mutants. Consistent with the patterns seen in roots, SA-sugar conjugates in shoots were more abundant in autophagy mutants, but this accumulation was only statistically significant for 2,3-DHBX in *atg11*. Overall these profiles indicate that many of the metabolic effects of autophagy disruption are already evident under control conditions, while some of them were enhanced by Pi limitation.

### Pi limitation had only a mild impact on root cytosolic protein degradation in mutant lines and on mitochondria abundance in *atg11*

To determine if protein degradation rates were similarly affected by Pi limitation, we compared protein abundance for 1045 proteins and degradation rates for 476 proteins among WT, *atg5,* and *atg11* roots under both control and Pi-limiting conditions. A principal component analysis (PCA) of these datasets showed that each genotype/treatment group could be separated by protein abundance and degradation rate (**Fig 8A-B**). Low Pi increased the abundance of vacuole proteins and decreased the abundance of Golgi proteins in WT; however, the same treatment caused an increase in vacuolar proteins in *atg11* but not in *atg5* whereas Golgi proteins were not significantly altered in either mutant (**Fig S7**). Low Pi did not induce significant changes in protein degradation rates in WT; however, it did decrease mitochondrial protein degradation rates in both *atg11* and *atg5,* and peroxisomal protein degradation rates in *atg5* (**Fig S8**).

**Fig 8.**
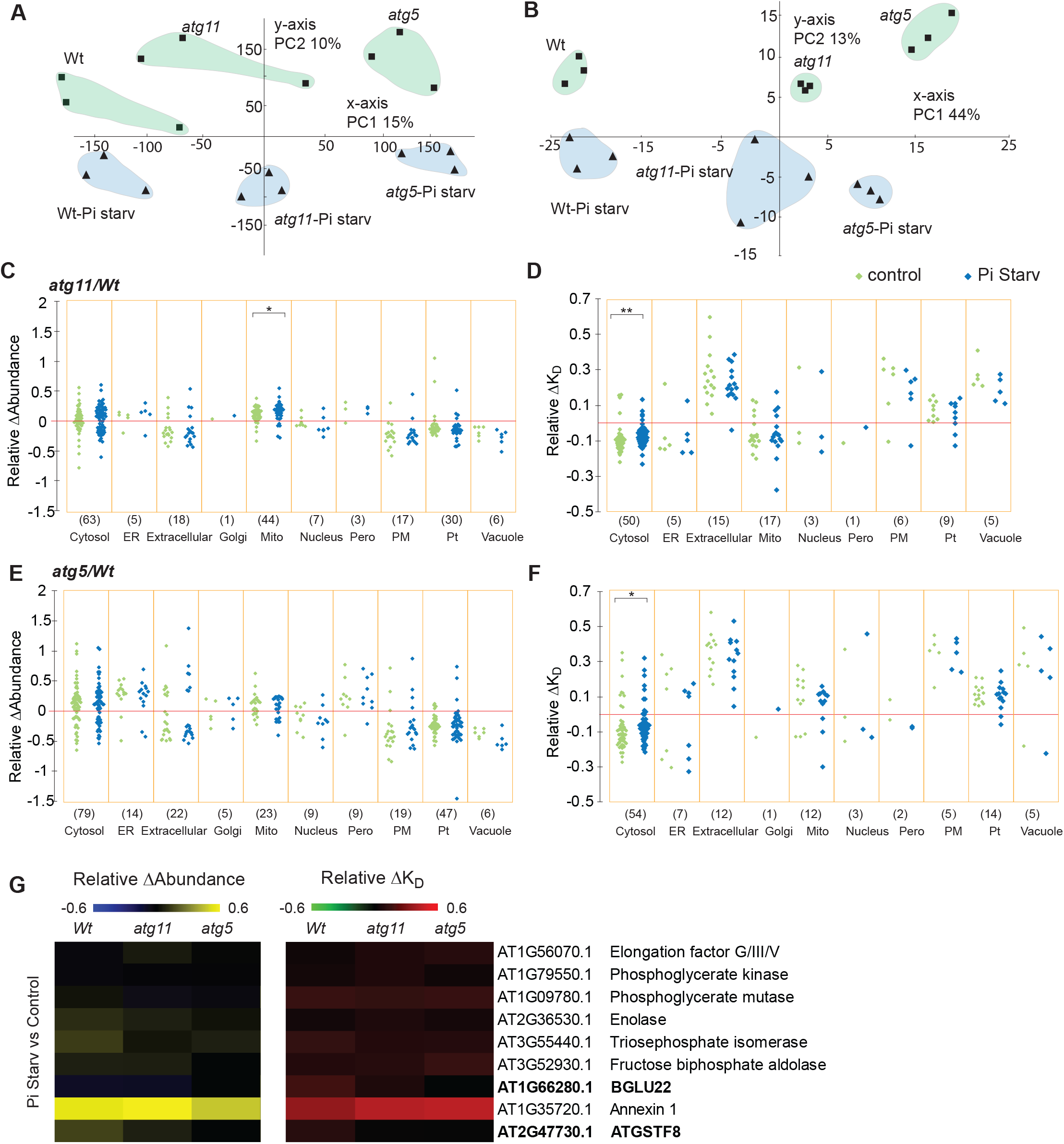
Pi limitation effects on changes of root protein abundance and degradation in *atg5* and *atg11*. (**A,B**) PCA analysis was applied to evaluate Pi limitation effects on protein abundance and degradation changes in *Wt*, *atg11* and *atg*5 using 1045 and 476 proteins, respectively. Protein abundance data were LN transformed before being used for PCA analysis. Principle components 1 and 2 (x and y axis) for all genotypes under both control and Pi starvation conditions are shown for protein abundance (**A**) and protein degradation (**B**). Relative changes of protein abundance and degradation between Wt and autophagy mutant lines were plotted to visualize Pi limitation effects on specific proteins of known location in root cells. Relative changes in protein abundance from 194 proteins in *atg11/Wt* comparisons (**C**) and from 233 proteins in a*tg5/Wt* comparisons (**E**) are shown as scattergrams. Relative changes in protein degradation rates from 111 proteins in *atg11/Wt* comparisons (**D**) and from 115 proteins in a*tg5/Wt* comparisons (**F**) are also shown as scattergrams. A nonparametric Kolmogorov-Smirno test was utilized for comparison of control and Pi starvation on distribution of relative changes in protein abundance and degradation rate of cellular localisations to evaluate the Pi limitation effect (** P<0.01, *P<0.05). Nine proteins show significantly faster degradation rates under Pi starvation conditions compared with control in Wt root. Relative degradation rate changes (relative ΔK_D_) and relative abundance changes (relative Δabundance) of these proteins in *Wt*, *atg5* and *atg11* between control and Pi limiting conditions are shown as heatmaps (**G**).

We then expressed the root datasets as relative changes in mutants and compared them between control and Pi-limiting conditions (**Fig 8C-F**). We found that mild Pi limitation further increased mitochondrial protein abundance in *atg11* compared to WT, but not in *atg5*. Unexpectedly, Pi limitation decreased the degree of differences in degradation rates of cytosolic proteins between the mutant lines and WT **(Fig 8D,F)**. This is seen in the narrower distribution of relative ΔK_D_ values under Pi limited conditions. However, we found nine proteins, including five cytosolic glycolytic enzymes, Annexin 1, the glutathione transferase ATGSTF8 and the ER-localized beta-glucosylase BGLU22, with significantly faster degradation rates under Pi limitation in WT (**Fig 8G, DataS10**). Intriguingly, the faster degradation of these proteins under low Pi did not lead to a decrease in their abundance; rather, four out of nine proteins were more abundant under low Pi conditions. This pattern is consistent with induced protein synthesis as a means to compensate for faster protein degradation under Pi limitation. Faster degradation of AtGSTF8 and BGLU22 under low Pi were only detected in WT but not in the autophagy mutants **(Fig 8G, DataS10)**.

### Pi limitation affects to degree of relative changes in abundance of chloroplast proteins in shoots and their degradation rates in both *atg5* and *agt11*

We also compared protein abundance of 782 proteins and degradation rates of 505 proteins among WT, *atg5,* and *atg11* shoots under control and Pi-limiting conditions. By applying a PCA, we found that protein abundance in WT samples could be separated from *atg5* and *atg11* under both control and Pi-limiting conditions, while *atg5* and *atg11* samples can be clearly separated under Pi limitation but not under control conditions (**Fig 9A**). In terms of protein degradation rates, WT samples could be fully separated by PCA from *atg5* and *atg11* under low Pi, but not under control conditions (**Fig 9B**). Pi limitation led to a decrease in cytosolic protein abundance in WT and *atg11* but not in *atg5.* Chloroplastic proteins accumulated in WT under low Pi whereas under similar conditions, chloroplastic protein abundances decreased in both mutant lines (**FigS7).** Pi limitation conditions altered protein degradation rates of chloroplastic proteins only in *atg5* (**FigS8).**

**Fig 9.**
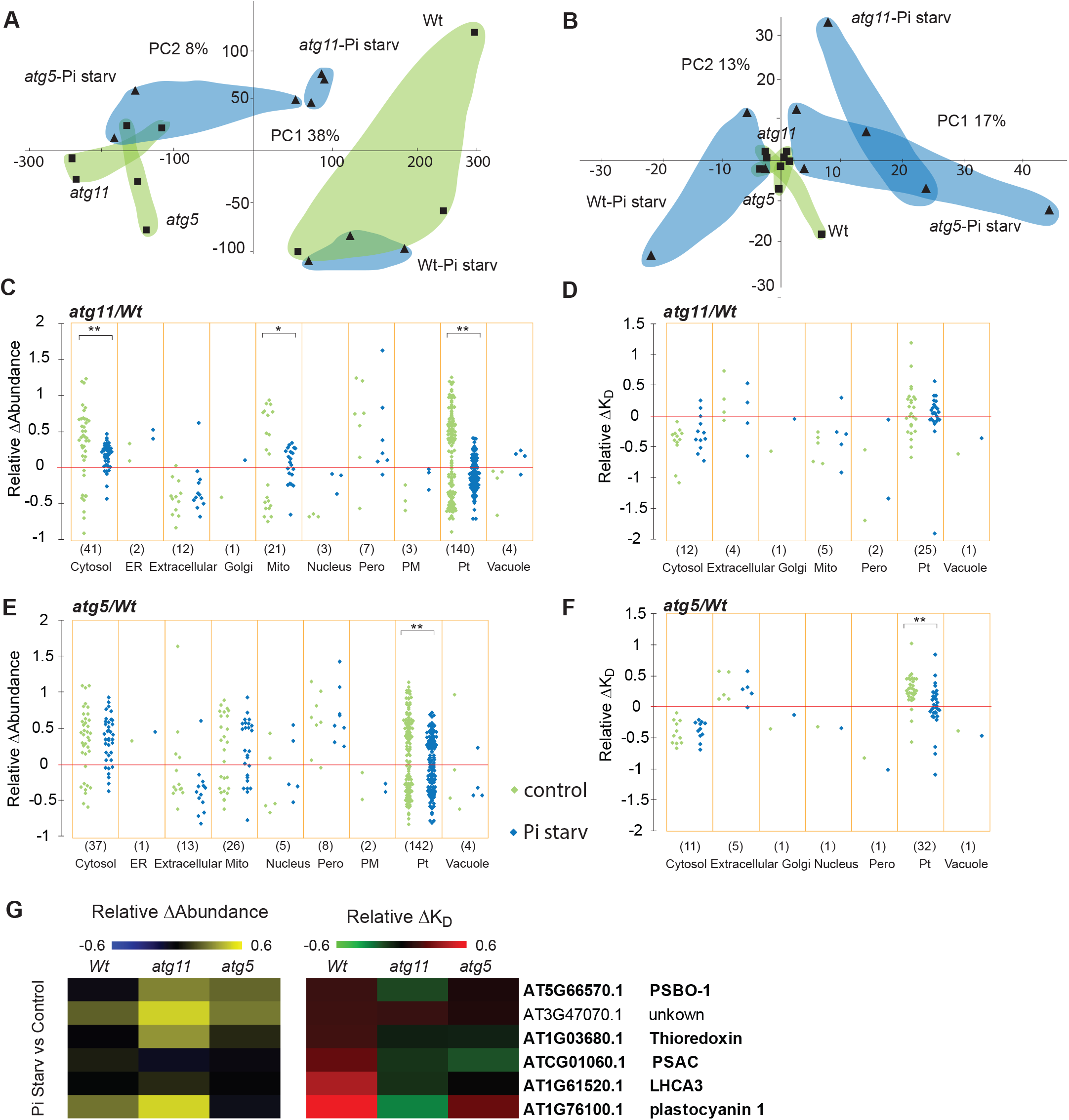
Pi limitation effects on changes of shoot protein abundance and degradation in *atg5* and *atg11*. (**A,B**) PCA analysis was applied to evaluate Pi limitation effects on protein abundance and degradation changes in *Wt*, *atg11* and *atg*5 using 782 and 505 proteins, respectively. Protein abundance data were LN transformed before being used for PCA analysis. Principle components 1 and 2 (x and y axis) for all genotypes under both control and Pi starvation conditions are shown for protein abundance (**A**) and protein degradation (**B**). Relative changes of protein abundance and degradation between Wt and autophagy mutant lines were plotted to visualize Pi limitation effects on specific proteins of known location in shoot cells. Relative changes in protein abundance from 238 proteins in *atg11/Wt* comparisons (**C**) and from 234 proteins in a*tg5/Wt* comparisons (**E**) are shown as scattergrams. Relative changes in protein degradation rates from 50 proteins in *atg11/Wt* comparisons (**D**) and from 52 proteins in a*tg5/Wt* comparisons (**F**) are also shown as scattergrams. A nonparametric Kolmogorov-Smirno test was utilized for comparison of control and Pi starvation on distribution of relative changes in protein abundance and degradation rate of cellular localisations to evaluate the Pi limitation effect (** P<0.01, *P<0.05). Six proteins show significantly faster degradation rates under Pi limiting conditions compared with control in Wt shoot. Their relative degradation rate changes (relative ΔK_D_) and relative abundance changes (relative Δabundance) in *Wt*, *atg5* and *atg11* between control and Pi limiting conditions are shown as heatmaps (**G**).

As done for the root datasets, we then expressed the shoot data as relative changes in mutants to facilitate comparisons between samples grown under control and Pi-limiting conditions (**Fig9C-F**). Low Pi again led to a narrower distribution of the abundance changes of chloroplastic proteins in both mutant lines compared to WT and smaller changes of protein degradation rate for chloroplast proteins in *atg5.* Conversely in *atg11*, Pi limitation was associated with a narrower distribution of changes in cytosolic and mitochondrial protein abundance without affecting the changes in protein degradation rate. Although there was no overall change in organellar degradation rate, six shoot proteins (PSBO-1, At3g47070, thylakoid phosphoprotein-At3g47070, thioredoxin-At1g03680, PSAC, LHCA3 and plastocyanin 1) showed significantly faster degradation rates under Pi limiting conditions in WT (**Fig 9G, Data S10**). Four of them, namely PSBO-1 (PSII), thioredoxin (chloroplast stroma), PSAC (PSI), and LHCA3 (PSI) showed unchanged or slower degradation rate in *atg11* and *atg5* under low Pi. These are therefore potential ATG11 and ATG5 associated targets of induced autophagy under low Pi. Plastocyanin 1 (thylakoid) showed a slower degradation rate and significant increase in abundance in *atg11* but faster degradation in *atg5* and non-significant change in abundance, which suggests its degradation is dependent on ATG11 but not ATG5. A notable exception to these trends was the thylakoid TSP9 phosphoprotein which is involved in photosystem state transition (Fristedt et al., 2009) that had an increased degradation rate but also accumulated in abundance in WT and mutant lines under low Pi, indicating its high protein synthesis rate and its turnover in all lines under Pi limitation.

## Discussion

### Investigation of protein turnover in autophagy mutant lines

Defects in autophagy can cause accumulation of autophagic protein cargo due to impaired degradation, but also lead to many changes in metabolite and transcript abundances, complicating the interpretation of cause and effect. Increase in transcript abundance may reveal up-regulation of gene expression (McLoughlin et al., 2018; McLoughlin et al., 2020), but that does not necessarily correlate with protein synthesis. In addition, estimating protein synthesis by ribosome profiling (Juntawong et al., 2014; Chotewutmontri and Barkan, 2016) or newly made protein labelling strategies (Wang et al., 2016) in autophagy mutants can be misleading since ribosomes themselves are targets of autophagy (Gretzmeier et al., 2017; McLoughlin et al., 2018), **Table S2**, **FigS4**). Focusing on steady-state protein abundance alone in impaired autophagy mutants also fails to identify proteins that may maintain homeostasis, either by using alternative degradative pathways or reducing protein synthesis. By focusing on protein degradation rates and correlating these with protein abundance in autophagy mutants, we circumvent some of these problems to reveal subsets of proteins that are directly influenced by autophagic processes, with or without compensatory changes in protein synthesis. Similar approaches in human fibroblasts (Zhang et al., 2016) and *Drosophila melanogaster* (Vincow et al., 2019) have also pinpointed specific protein complexes and organelles that are differentially affected by autophagy in other organisms.

### Common changes in cytosolic protein abundance and degradation rates in *atg11* and *atg5*

The *ATG11* and *ATG5* mutants used in this report were previously shown to be *bona fide* single gene mutants in Arabidopsis, to have impaired autophagic fluxes detected by vacuolar delivery of *ATG8-GFP,* and to share typical early senescence phenotypes at late developmental stage and under nutrient limitation conditions (Thompson et al., 2005; Yoshimoto et al., 2009b; Li et al., 2014; Li and Vierstra, 2014). We show *atg11* and *atg5* share many common differences in cytosolic and organellar protein abundances and associated changes in protein degradation rates, although *atg5* typically showed larger relative changes in protein abundance (**Fig 1&4, Fig S4&S5**). The larger differences in *atg5* are broadly consistent with the severity of this mutant’s senescence phenotype compared with *atg11* (Yoshimoto et al., 2009b; Li et al., 2014), and the established roles of ATG5 and ATG11 in the autophagy process; namely ATG5 acting in the core conjugation cascade and ATG11 acting in a regulatory complex.

Glycolytic enzymes including phosphoglycerate kinase, enolase, triosephosphate isomerase and fructose bisphosphate aldolase showed slower protein degradation rates in both *atg11* and *atg5*, but these led to only a mild increase in abundance of these enzymes (0-24%) in autophagy mutants. Interactions between autophagy and glycolytic enzymes have been previously reported in plants and other organisms and can be complex (Han et al., 2015; Henry et al., 2015; Watson et al., 2015; Qian et al., 2017a; Qian et al., 2017b). Firstly, glycolytic enzymes play roles in autophagic flux regulation. For example, glyceraldehyde-3-phosphate dehydrogenases (GAPDHs) negatively regulate autophagy in Arabidopsis (Henry et al., 2015) and in tobacco GAPDHs can reduce autophagy activities by binding to ATG3 (Han et al., 2015). In contrast, phosphoglycerate kinase 1 (PGK1) can induce autophagy under cellular stress conditions in mammals through phosphorylating of Beclin1 (Qian et al., 2017b). Secondly, autophagy can downregulate glycolysis metabolism through selective degradation of enzymes. For example, selective degradation of hexokinase (HK) in human liver cancer cells during autophagy (Jiao et al., 2017). Here we show the basal rate of degradation of glycolytic enzymes in Arabidopsis is partially due to autophagy, but impaired autophagy may be compensated for by changes in glycolytic enzyme synthesis that prevent their accumulation. Recently, proximity-dependent biotinylation screening of *in vivo* interactions confirmed GAPDH and fructose bisphosphate aldolase are bound by ATG8 in plants (Macharia et al., 2019).

The Chaperonin Containing T-complex polypeptide-1 (CCT) protein complex in human cell lines is present in immunopurified autophagosomes, degrades slowly in autophagy mutants (Dengjel et al., 2012; Zhang et al., 2016), and can restrict neuropathogenic protein aggregation via autophagy in human cell lines and fruit fly (Pavel et al., 2016). Although well documented in animals, a 20S protein complex consisting of eight CCT subunits was only recently reported in plants (Ahn et al., 2019; McWhite et al., 2020). In this study, we found five CCT protein complex subunits (CCT1-4, 8) increased in abundance and had slower rates of degradation in roots of both autophagy mutants **(Table S1, Fig S4),** supporting the hypothesis that CCT is an autophagy target in Arabidopsis roots. CCT is known to affect the folding and stability of tubulin in Arabidopsis and mutants with deficient CCT function show depletion of cortical microtubules and reduced alpha and beta tubulin abundance due to increased degradation (Ahn et al., 2019). Interestingly, we found that five beta tubulins (TUB2,4,6,8,9) but not alpha tubulins, showed decreased abundance in *atg11* and *atg5*. It is unclear whether the decreased abundance of beta tubulin is a direct or indirect effect of impaired autophagy, but microtubules are important for autophagy. Microtubules can interact with autophagic proteins and play roles in pre-autophagosome structure, autophagy induction, formation and movement (Mackeh et al., 2013). In plants, microtubules are proposed to aid autophagosome delivery to the vacuole with the help of FYVE and coiled-coil domain–containing (FYCO) proteins that bind ATG8 and PI3K on the autophagosome outer membrane (Marshall and Vierstra, 2018). We also found actin (ACT7) and the actin-interacting proteins VILIN4 and PROFILIN1/2 showed decreased abundance in both mutants. In yeast, actin filaments are only involved in selective but not bulk autophagy (Hamasaki et al., 2005; Reggiori et al., 2005; Monastyrska et al., 2009). In mammals, actin was found to be required for both selective and bulk autophagic degradation (Kast and Dominguez, 2017; Xu et al., 2018). In plants, actin filaments seem to be dispensable for bulk autophagy in tobacco (Zheng et al., 2019), but it is unclear whether actin is needed for any form of selective autophagy in plants. Our results thus suggest that the homeostasis of the plant CCT complex is controlled by autophagic degradation and that accumulation of CCT complex subunits correlated with decreases in the abundance of components of the cytoskeleton, which warrants further investigation.

### Changes in organelle abundance and degradation rates in *atg11* and *atg5*

Many proteins with decreased degradation rates in both root and shoot tissues of autophagy mutants localize to the ER, peroxisomes, or mitochondria (**Fig 5**). This supports the notion that ER, peroxisome and mitochondrial proteins are autophagy cargo in both photosynthetic and non-photosynthetic tissues. Interestingly, we found that in roots, a high proportion of these proteins increased in abundance, while in shoots a high proportion decreased in abundance **(Fig 1)**. This means that proteins with slower degradation showed reduced abundance in shoot but accumulated in roots, suggesting that photosynthetic tissues have more plasticity for transcription and translational control during autophagy than root tissues. The more prominent deployment of alternative proteostasis/protein recycling mechanisms in shoots than in roots is also consistent with the more drastic decrease in amino acids levels in roots than in shoots of autophagy mutants (**Fig 7**).

In roots, organelle proteins with faster degradation rates in autophagy mutants were predominantly localized to plastids, apoplast, plasma membrane, and vacuoles. Whereas we did not find evidence of increases in plastid protease abundance in roots (**Data S1**), other degradative pathways can deliver portions of plastids to the vacuole (Izumi et al., 2017; Otegui, 2018; Zhuang and Jiang, 2019). Therefore, it is possible that ATG5/ATG11-independent pathways that mediate plastid turnover are stimulated in roots of autophagy-deficient mutants. The content of extracellular, plasma membrane, and vacuolar proteins is closely associated with the rate of intracellular vesicle trafficking. Proteins reach the vacuole through the secretory and endocytic/endosomal pathways as well as through autophagy (Marty, 1999; Pereira et al., 2014; Zhang et al., 2014; Shimada et al., 2018). Deficient autophagy in *atg5* and *atg11* correlated with a decreased abundance of both tonoplast and vacuolar lumen proteins (**Fig S4, Fig S5**). The outer membrane of the autophagosome is integrated into the tonoplast upon fusion and it is therefore assumed to supply large quantities of membrane to vacuoles. Consistently, although not statistically significant, we noticed by electron microscopy a consistent decrease in tonoplast membrane in actively vacuolating cells of both *atg5* and *atg11* root cells. We also found that CLATHRIN HEAVY CHAIN 1 (CHC1, At3g11130) was more abundant in roots of *atg5* (**Fig S4**). Clathrin is associated with endocytosis at the plasma membrane and sorting at the TGN and endosomes (Gao et al., 2019). The altered abundance of clathrin and other trafficking components could alter both endocytosis/endosomal and exocytosis rates, contributing to the fast turnover and low abundance for both plasma membrane and extracellular proteins seen here in autophagy mutants. However, future experimental evidence is still needed to investigate the endocytosis/exocytosis processes in *atg5* and other autophagy mutants.

### Protein abundance and degradation rate specific changes in *atg11* and *atg5*

The most severe molecular alterations in *atg5* were the five-fold increases in specific ER-resident proteins. This differential effect on ER homeostasis in *atg5* also correlated with accumulation of vesicle transport-associated proteins, such as three COP1 coatomers (alpha, delta and gamma subunits) in *atg5*. COP1 is essential for retrieval of proteins with di-lysine motifs from Golgi stacks back to the ER (Wang et al., 2018), intra Golgi transport, and Golgi maintenance. Interestingly, we observed in *atg5* but not in *atg11* root cells abnormal membranous structures with assembled coats of unknown nature reminiscent of Golgi and/or ER membranes **(Fig 3)**. Whereas the origin of these abnormal, coated, membranous structures in *atg5* cells is unknown, the mis-regulation of COP1 components in mammalian cells induces the re-localization of Golgi, TGN, and ER markers into large membranous structures (Styers et al., 2008). The fungal toxin brefeldin A inhibits the assembly of the COP1 coat and also results in large abnormal membranous bodies (BFA bodies) in plants, that contain Golgi, TGN, and endosomal proteins (Nebenfuhr et al., 2002; Lam et al., 2009; Berson et al., 2014). In addition, the loss of COP1 subunits also leads to the accumulation of abnormal autophagosomes not fully capable to fusing with lysosomes (Razi et al., 2009).

A higher proportion of mitochondrial proteins with slower degradation rates was found in *atg11* compared with *atg5,* in both root and shoot tissues (**Table S1, S2**). Higher abundance of mitochondrial proteins was also common in both shoot and root of *atg11,* but only in shoots of *atg5* **(Fig 1)**. ATG11 has been reported to be essential for senescence-induced mitophagy in Arabidopsis photosynthetic tissues (Li et al., 2014; Li and Vierstra, 2014) and ATG5-dependent mitophagy has been recently reported in Arabidopsis cotyledons and roots (Ma et al 2021); however, *in vivo* changes in degradation rate of specific mitochondrial proteins in either *atg11* or *atg5* under control conditions has not been reported previously to our knowledge. Interestingly, although chloroplast proteins show general increases in abundance in shoots of both *atg11* and *atg5*, we only found a higher proportion of chloroplast proteins with slower degradation rates in shoots of *atg11* (**Fig 2**, **Table S2**). These same chloroplast proteins showed faster protein turnover rates in *atg5*. We interpret this to mean that chloroplast proteins accumulated in *atg11* through deficient degradation but through enhanced synthesis in *atg5*. Taken together, the different patterns of degradation and abundance changes in mitochondrial and chloroplastic proteins in the mutant lines support a specialized role of ATG11 in basal level mitochondrial and chloroplast protein homeostasis and highlight specific organelle proteins that are good indicators of this role.

### Pi limitation caused autophagy-dependent cytosolic protein degradation and vacuole biogenesis in roots, and chloroplast degradation in shoots

Pi limitation has been reported to induce autophagy in yeast and plants (Tasaki et al., 2014; Yokota et al., 2017; Naumann et al., 2019) and our data indicate we could reproduce this effect in hydroponically-grown Arabidopsis plants (**Fig 6**). Pi limitation-induced autophagy is reported to contribute to vacuole biogenesis (Gao et al., 2017), which is consistent with the general increase in abundance of vacuolar proteins in roots of WT plants grown in Pi-limiting conditions (**Fig S7**). A similar increase in vacuolar proteins was observed in roots of *atg11* but not of *atg5*, suggesting ATG11 is not essential for autophagy-dependent vacuole biogenesis under Pi-limiting conditions. A new ATG1-independent autophagy mechanism in prolonged carbon starvation conditions was recently reported (Huang et al., 2019), so this might explain the activation of an autophagic pathway independent of the ATG1 kinase complex in *atg11*. However, the role of ATG11 in mitochondrial degradation was not diminished under Pi limitation (**Fig 8C**), indicating that ATG1-independent autophagy in *atg11* cannot compensate for deficient mitochondria degradation. We also found that Pi limitation attenuated the cytosolic protein abundance differences observed between WT and autophagy mutants, without affecting protein degradation rates. This could indicate complementary transcription/translation changes induced by Pi limitation in these autophagy mutants.

In roots, BGLU22 and ATGSTF8 had faster turnover rates under Pi limitation in WT but not in either autophagy mutant (**Fig 8G**). While BGLU22 showed a slight abundance reduction, ATGSTF8 accumulated in WT. This indicates that in WT plants growing in Pi-limiting conditions, BGLU22 turnover is associated with the stimulated autophagic degradation while ATGSTF8 increased turnover is compensated with even greater protein synthesis. BGLU22 localizes to root ER bodies is an EE-type myrosinase that can break down aliphatic glucosinolates during stress conditions (Sugiyama and Hirai, 2019). The faster degradation of BGLU22 under Pi-starvation suggest that ER bodies containing BGLU22 may be delivered to the vacuoles containing the aliphatic glucosinolates by autophagy.

All six proteins that degraded faster under Pi limiting conditions in WT shoots were chloroplast-resident proteins. Four of them showed unchanged or slower degradation rate in *atg11* and *atg5* **(Fig 9G),** suggesting a role for Pi limitation in ATG11/5 associated autophagic chloroplast degradation. These unchanged or slower degrading proteins were components of photosystems and their light harvesting complexes (PSBO-1, PSAC, LHCA3), and a plastid thioredoxin (thioredoxin M1) that regulates photosynthetic acclimation in fluctuating light intensities by regulating the export of excess reductive power from the chloroplasts (Thormahlen et al., 2017). An electron carrier between photosystems (plastocyanin 1) showed slower degradation and a significant increase in abundance in *atg11* but faster degradation rate in *atg5* and no change in abundance, which suggest its degradation is dependent on ATG11 but not ATG5 (**Data S10**). These represent useful protein markers to study general or selective chloroplast degradation by autophagy.

## Methods

### Arabidopsis hydroponic plants preparation and ^15^N labelling

*Arabidopsis thaliana* accession Columbia-0 (WT), *atg5* and *atg11* plants were grown under 16/8-h light/dark conditions with cool white T8 tubular fluorescent lamps 4000K 3350 lm (Osram, Germany) with intensity of 100–125 μmol m^−2^ s^−1^ at 22 °C. The hydroponic protocol was as described previously (Waters et al., 2012) and used a modified Hoagland solution (2 mM CaCl_2_, 6 mM KNO_3_, 0.5 mM NH_4_NO_3_, 0.5 mM MgSO_4_, 0.25 mM KH_2_PO_4_, 0.05 mM KCl and 0.04 mM Fe-EDTA) supplemented with micro elements (25 μM H_3_BO_3_, 2 μM MnCl_2_, 2 μM ZnSO_4_, 0.5 μM CuSO_4_, 0.15 μM CoCl_2_ and 0.25 μM (NH_4_)_6_Mo_7_O_24_) and 2.6 mM MES, and the pH was adjusted to 5.8-6.0. Seeds of different lines (WT, *atg5* and *atg11*) were planted on the growth hole of agar stuffed in lids of 1.5-ml black tubes sitting in 24-well floater tubes racks containing 160 ml growth medium. The seeds were vernalized under 4 °C for 2-3 days before being transferred to the growth chambers. Half-strength growth medium was used for the first week. A single plant was placed in every tube lid and four tubes lids in each floater tube rack (**FigS1**). The growth medium was changed every 5 days. Unlabelled Arabidopsis plants were grown for 21 days until they reached leaf production stage 1.10 (T0) (Boyes et al., 2001) in natural abundance medium as noted above. To obtain a fully labeled ^15^N protein reference standard, ^15^N medium (with 6 mM K^15^NO_3_, 0.5 mM ^15^NH_4_^15^NO_3_) was used to replace the natural abundance nitrogen in the medium and plants were grown from seed in this medium for 26 days. For progressive ^15^N labelling, the growth medium was discarded and the growth racks rinsed four times with fresh medium without nitrogen (no KNO_3_ or NH_4_NO_3_) to ensure the old solution was washed out. A total of 160 ml of ^15^N medium (6 mM K^15^NO_3_, 0.5 mM ^15^NH_4_^15^NO_3_) was added for every four plants and the plants were grown for three days before collecting leaf and root tissues for separate total protein extraction (**FigS9**). Root/shoot from two plants in one rack were pooled as a biological replicate, three biological replicates were collected.

### Protein extraction, in-solution digestion, high pH HPLC separation and LC-MS analysis of *tryptic peptides*

The root/shoot samples (0.1-0.2 g) from fully ^15^N labelled reference, ^15^N progressively labeled and unlabeled of three lines (WT, *atg5* and *atg11*) were snap frozen in liquid nitrogen and homogenized using Qiagen tissue lysis beads (5 mm) by vortex. A total plant protein extraction kit (PE0230-1KT, Sigma Chemicals) was used to extract root/shoot total proteins. The final pellet of total protein was dissolved in Solution 4 and then reduced and alkylated by tributylphosphine (TBP) and iodoacetamide (IAA) as described in the Sigma manual. The suspension was centrifuged at 16,000 g for 30 min and the supernatant was assay for protein concentration by amido black quantification as described previously (Liu et al., 2012a).

A total of 100 μg root/shoot proteins from progressively ^15^N labelled samples were digested in solution as described previously (Nelson et al., 2014). A total of 50 μg of unlabeled root/leaf protein samples noted above was mixed individually with 50 μg of the fully ^15^N-labelled reference and digested in solution by trypsin. Each sample was separated into 96 fractions by high pH HPLC separation and further pooled into 12 fractions and each fraction was analyzed by mass spectrometry. Filtered samples (5 μl each) were loaded onto a C18 high-capacity nano LC chip (Agilent Technologies) using a 1200 series capillary pump (Agilent Technologies) as described previously (Li et al., 2017).

### MS data analysis, calculations of *K_D_* and relative abundance values

Agilent .d files were converted to mzML using the Msconvert package (version 2.2.2973) from the Proteowizard project, and mzML files were subsequently converted to Mascot generic files using the mzxml2 search tool from the TPPL version 4.6.2. Mascot generic file peak lists were searched against an in-house *Arabidopsis* database comprising ATH1.pep (release 10) from The *Arabidopsis* Information Resource (TAIR) and the *Arabidopsis* mitochondrial and plastid protein sets (33621 sequences; 13487170 residues) (Lamesch et al., 2012), using the Mascot search engine version 2.3 and utilizing error tolerances of 100 ppm for MS and 0.5 Da for MS/MS; “Max Missed Cleavages” set to 1; variable modifications of oxidation (Met) and carbamidomethyl (Cys). We used iProphet and ProteinProphet from the Trans Proteomic Pipeline (TPP) to analyze peptide and protein probability and global false discovery rate (FDR) (Nesvizhskii et al., 2003; Deutsch et al., 2010; Shteynberg et al., 2011). The reported peptide lists with p=0.8 have FDRs of <3% and protein lists with p=0.95 have FDRs of <0.5%. Quantification of LPFs (labeled peptide fraction) were accomplished by an in-house R script which was written originally in Mathematica (Nelson et al., 2014). A median polish method described previously was used for data analysis (Li et al., 2017). Measured protein degradation rate 0.1 d^−1^ was used to calculate the FCP (fold change protein) for shoot samples. For root tissues, measured FCP based on fresh weight was measured before and after progressive ^15^N labelling. A measured degradation rate 0.5 d^−1^ was determined and then applied to calculate FCP in samples of *Wt* and mutant lines, which were applied for degradation rate calculations. We determined changes in specific protein abundance using a fully labeled ^15^N protein reference standard. Protein abundance was represented as ratio to reference and normalized to all samples (three lines under both control and Pi starvation conditions) as previously reported (Li et al., 2017). Relative ΔAbundance (i.e. (mutant–Wt)/Average (mutant and Wt)) was used to describe the level of changes between mutant vs Wt or treatment vs control.

### RNA extraction and Q-PCR analysis

We collected leaf 5 from all three lines at 21 days, and collected leaf 6 after three days Pi starvation treatment. The shoot samples (~0.1g) from three lines (*Wt*, *atg5* and *atg11*) under control/Pi starvation conditions were snap frozen in liquid nitrogen and homogenized to powder using Qiagen tissue lysis beads (2 mm) by a homogenizer. RNA was extracted using Spectrum^TM^ Plant Total RNA kit (Sigma-Aldrich, STRN250-1KT)with On-Column DNase treatment (Sigma-Aldrich, DNASE70) following manufacturer’s instructions. 500ng of RNA was used for cDNA synthesis with iScript cDNA synthesis kit (Bio-rad, 1708890). Transcripts of *spx1 (F- TGCCGCCTCTACAGTTAAATGGC*, *R-TGGCTTCTTGCTCCAACAATGG*), *atg8h (F-TGCAGTTAGATCCATCCAAAGCTC*, *R-TCCATGCGACTAGCGGTTTGAG*) and *atg7 (F-ACGTGGTTGCACCTCAGGATTC*, *R- ACTAAGAGTTCAACGGCGAGAGC*) were quantified using QuantiNova SYBR green PCR kit (Qiagen, 208056) with LightCycler380 in *Wt*, *atg5/11* lines under both control and Pi starvation conditions. We did four biological replicates for most samples except *atg11* post starvation treatment. QPCR data were normalized to housekeeping genes AKT2 *(F- GGTAACATTGTGCTCAGTGGTGG*, *R-AACGACCTTAATCTTCATGCTGC*) and UBQ10*(F-CTGCGACTCAGGGAATCTTCTA*, *R-TTGTGCCATTGAATTGAACCC*) before analysed using geometric averaging of multiple control genes (Vandesompele et al., 2002; Czechowski et al., 2005) before being compared.

### Pi concentration measurement by a colorimetric assay

*Wt*, *atg5* and *atg11* lines were grown hydroponically till leaf production stage 1.10 (T0). Growth containers were rinsed with water for complete phosphate depletion. For Pi starvation treatment, plant growth media was replaced with Hoagland solution without phosphate and grown for three days (T3-Pi starvation). Hoagland solution with phosphate was used for control plants (T3). Inorganic concentration in root/shoot tissues of three lines were measured by a colorimetric assay. Inorganic phosphate was extracted in 500 μl water from 10-mg frozen powdered samples. The concentration of P_i_ was determined spectrophotometrically at 820 nm after a 90-min reaction at 37°C in the presence of 1.4% w/v ascorbate and 0.36% w/v ammonium molybdate in 1 N H_2_SO_4_ (Ames, 1966).

### Maximum quantum yield of PSII measurement by IMAGING-PAM

Leaf production stage 1.10 Arabidopsis plants (T0-grown in hydroponics for ~21 days post-germination) were washed by Hoagland media without phosphate and then grown for another three days in fresh growth media (T3-control) or growth media without phosphate (T3-phosphate starvation). Whole plants were dark adapted least 20 mins before being measured by a MAXI version of the IMAGING-PAM. A color gradient was used to demonstrate the Fv/Fm (maximum quantum yield of PSII) values which were measured by IMAGING-PAM in leaves of the whole rosette. One biological replicate was a combination of measured Fv/Fm values in six leaves in two Arabidopsis plants.

### Confocal laser scanning microscopy

GFP-ATG8a plants were grown hydroponically till leaf production stage 1.10. Whole plants were transferred into normal or −Pi growth media and grown for another three days. E64d was supplemented into growth media 24 hours before the confocal laser scanning microscopy experiment to a final concentration of 100 μM. A Nikon A1Si confocal microscope equipped with laser line 488-nm excitation and emission band-pass filter of 500-520 nm, and controlled by a NIS element AR software package (version 4.13.01, Build 916) was used. Images were acquired using a 20x lens (Nikon CFI Plan Apo VC 20× 0.75 N.A.) with pinhole diameter of 2.5 airy units (corresponds to the optical slice of 4.37 um). Autophagic puncta (AP) of representative images were counted by the ‘Analyze Particles’ function of ImageJ. AP numbers in each Z-stack were plotted. The distribution of AP number under control and Pi limitation conditions in the representative image were compared by Kolmogorov-Smirnov test for significance.

### Transmission electron microscopy

Arabidopsis seedlings were grown for 24 days in hydroponic conditions as described above. Root tips were excised and placed in freezing planchettes containing 0.1M sucrose and high-pressure frozen in a Baltec HPM 010. Samples were high-pressure frozen in 2% (w/v) OsO_4_ in anhydrous acetone in dry ice overnight and warmed to room temperature on a rocker with slow agitation for several hours, until they reached at room temperature. After several acetone rinses and the planchets removed, samples were infiltrated in a series of Epon resin changes polymerizing at 60°C for 24 h. Sections were stained with 2% uranyl acetate and lead citrate (2.6% lead nitrate and 3.5% sodium citrate, pH 12) and imaged in a Philips CM120 transmission electron microscope. Morphological measurements were done using FIJI (Schindelin et al., 2012).

### Metabolite Extraction

Plant tissues (15–50 mg) were collected at specified time points and immediately snap-frozen in liquid nitrogen. Samples were ground to fine powder and 500 μl of cold metabolite extraction solution (90% [v/v] methanol, spiked with 2 mg/ml ribitol, 6 mg/ml adipic acid, and 2 mg/ml and ^13^C-leucine as internal standards). Samples were immediately vortexed and shaken at 1,400 rpm for 20 min at 75°C. Cell debris was removed by centrifugation at 20,000 × g for 5 minutes. For each sample, 100 or 400 μl of supernatant was transferred to a new tube and either proceeded to derivatization for LC-MS analysis or dried using a SpeedVac.

### Analyses of salicylic acid, organic acids and amino acids by selective reaction monitoring using triple quadrupole (QQQ) mass spectrometry

For LC-MS analysis of organic acids, sample derivatization was carried out based on previously published methods with modifications (Han et al., 2013). Briefly, for each of 100 μL of sample, 50 μL of 250 mM 3-nitrophenylhydrazine in 50% methanol, 50 μL of 150 mM 1-ethyl-3-(3-dimethylaminopropyl) carbodiimide in methanol, and 50 μL of 7.5% pyridine in 75% methanol were mixed and allowed to react on ice for 60 minutes. To terminate the reaction, 50 μL of 2 mg/mL butylated-hydroxytoluene in methanol was added, followed by the addition of 700 μL of water. Derivatized organic acids were separated on a Phenomenex Kinetex XB-C18 column (50 × 2.1mm, 5μm particle size) using 0.1% formic acid in water (solvent A) and methanol with 0.1% formic acid (solvent B) as the mobile phase. The elution gradient was 18% B at 1 min, 90% B at 10 min, 100% B at 11 min, 100% B at 12 min, 18% B at 13 min and 18% B at 20 min. The column flow rate was 0.3 mL/min and the column temperature was maintained at 40 °C. The QQQ-MS was operated in the negative ion mode with multiple reaction monitoring (MRM) mode.

For measuring salicylic acid and amino acids, dried samples were resuspended in 100 μL HPLC-grade water before they were filtered to remove insoluble debris. Metabolites were separated on an Agilent Poroshell 120 Bonus-BP column (100 × 2.1 mm, 2.7μm internal diameter) using 0.1% formic acid in water (solvent A) and acetonitrile with 0.1% formic acid (solvent B) as the mobile phase. For the analysis of amino acids and sugars, the elution gradient was 0% B at 1 min, 1% B at 4 min, 10% B at 6 min, 100% B at 6.5 min, 100% B at 8 min, 0% B at 8.5 min and 0% B at 15 min. The column flow rate was 0.25 mL/min, the column temperature was kept at 40 °C. The QQQ-MS was operated in the positive ion mode with MRM mode. For salicylic acid, the elution gradient was 0% B at 1 min, 1% B at 3 min, 95% B at 23 min, 100% B at 23.2 min, 100% B at 25 min, 0% B at 25.5 min and 0% B at 34 min. The column flow rate was 0.20 mL/min and the column temperature was set to 40 °C. The LC-MS was operated in the negative ion mode with MRM mode.

A 0.5 μL or a 15 μL aliquot of each sample were injected and analysed by an Agilent 1100 HPLC system coupled to an Agilent 6430 Triple Quadrupole (QQQ) mass spectrometer equipped with an electrospray ion source. Data acquisition and LC-MS control were done using the Agilent MassHunter Data Acquisition software (version B06.00 Build 6.0.6025.4). The autosampler was kept at 10°C. The QQQ-MS was operated in MRM mode using the following operation settings: capillary voltage, 4000V; drying N_2_ gas and temperature, 11 L/min and 125 °C respectively; Nebulizer, 15 psi. All optimised MRM transitions for each target were listed in **Data S12**. All data was analysed using MassHunter Quantitative Analysis Software (version B.07.01, Build 7.1.524.0). Metabolites were quantified by comparing the integrated peak area with a calibration curve obtained using authentic standards, and normalised against fresh weight and internal standards.

### Measurement and identification of sugars and secondary metabolites by quadrupole/time-of-flight mass (Q-TOF) spectrometry

Analyses of sugars and secondary metabolites were performed using an Agilent 1100 HPLC system coupled to an Agilent 6510 Quadrupole/Time-of-Flight (Q-TOF) mass spectrometer equipped with an electrospray ion source. Data acquisition and LC-MS control were carried out using the Agilent MassHunter Data Acquisition software (version B02.00). Separation of metabolites was performed using a Luna C18 column (Phenomenex; 150 × 2 mm, 3 μm particle size). The mobile phase consisted of 97:3 water:methanol with 10 mM tributylamine and 15 mM acetic acid (solvent A) and 100% methanol (solvent B). The gradient program was 0% B 0 min, 1% B 5 min, 5% B 15 min, 10% B 22 min, 15% B 23 min, 24% B 25 min, 29% B 80 min, 95% B 81 min, 95% B 82 min, 0% B 83 min and 0% B 97min. The flow rate was 0.2 mL/min, with column temperature kept at 35°C and samples at 10°C. The Q-TOF was operated in MS mode with negative ion polarity using the following operation settings: capillary voltage, 4000V; drying N_2_ gas and temperature, 10 L/min and 250 °C respectively; Nebulizer, 30 psi. Fragmentor, skimmer and octopole radio frequency (Oct1 RF Vpp) voltages were set to 110V, 65V and 750V respectively. The scan range was 70-1200 m/z and spectra were collected at 4.4 spectra/s which corresponded to 2148 transients/spectrum. All MS scan data was analysed using MassHunter Quantitative Analysis Software (version B.07.01, Build 7.1.524.0). Peaks were normalised against sample weight and the internal standard. For identification of metabolites without authentic standards, Q-TOF was operated in Targeted MS/MS mode with negative ion polarity using the same MS settings as outlined above. The MS/MS scan range was 40-1000 m/z and spectra were collected at 3.7 spectra/s which corresponded to 2603 transients/spectra. For each metabolite target, the retention time window was set to ±1 min, isolation width was set to narrow (~1.3 m/z), 10− to 20− and 40-eV collision energies were used and the acquisition time was set to 180 ms/spectra. The identity of each unknown was verified by comparing MS/MS fragment ions with published data (Stobiecki et al., 2006; Lee et al., 2008; Rochfort et al., 2008; Matsuda et al., 2009; Bartsch et al., 2010; Bialecki et al., 2010; Zhang et al., 2013; Lin et al., 2014; Hohner et al., 2018). The expected m/z, retention time and the method for identification were listed in **DataS12**.

## Open accessible data

PRIDE Project Name: To investigate the role of autophagy in Arabidopsis root cellular protein turnover and proteostasis (15N Spike-in root)

Project accession: PXD010992

PRIDE Project Name: To investigate the role of autophagy in Arabidopsis shoot cellular protein turnover and proteostasis (15N Spike-in shoot)

Project accession: PXD010948

PRIDE Project Name: To investigate the role of autophagy in Arabidopsis root cellular protein turnover and proteostasis

Project accession: PXD010900

PRIDE Project Name: To investigate the role of autophagy in Arabidopsis shoot cellular protein turnover and proteostasis

Project accession: PXD010932

## Acknowledgements

*atg5-1* (SAIL_129_B07), *atg11-1 (*SAIL_1166_G10) and the *GFP-ATG8a* line were kindly provided by Professor Richard Vierstra, Washington University, St Louis, MO. We thank Julio Paez-Valencia for his assistance growing plants for transmission electron microscopy analysis and Zhirui Chen for her assistance for confocal image quantifications.

## Author Contribution

LL and AHM designed the research; CPL, AWY and MB performed plant culture and biochemical experiments; MSO performed and analyzed TEM data. Mass spectrometry and analysis was performed by LL. LL, AHM and MSO contributing to the writing and revision of the article.

## Supplemental Tables

**Table S1** Identification of putative ATG5 and ATG11 targets in Arabidopsis roots as proteins with higher abundance and slower degradation rates in *atg5 or atg11* compared with wild-type.

**Table S2** Identification of putative ATG5 and ATG11 targets in Arabidopsis shoots as proteins with higher abundance and slower degradation rates in *atg5 or atg11* compared with wild-type.

## Supplemental Figures

**Fig S1** Arabidopsis *atg5* and *atg11* phenotypes compared to Wt plants.

**Fig S2** Changes in protein abundance in roots and shoots of Arabidopsis autophagy mutants.

**Fig S3** Significant changes in abundance of ribosome and proteasome subunits in Arabidopsis autophagy mutants.

**Fig S4** Significant changes in relative Δabundance of 241 root proteins in Arabidopsis autophagy mutants.

**Fig S5** Significant changes in relative Δabundance of 265 shoot proteins in Arabidopsis autophagy mutants.

**Fig S6** Changes in protein degradation rate (K_D_) in roots and shoots of Arabidopsis autophagy mutants compared to Wt.

**Fig S7** Effects of Pi limitation on protein abundance in roots and shoots of Wt and autophagy mutants.

**Fig S8** Effects of Pi limitation on protein degradation rates in roots and shoots of Wt and autophagy mutants.

**Fig S9** Workflow of analysis to determine protein abundance, protein degradation rates and metabolite abundances in samples from Arabidopsis plant tissues.

## Supplemental Data

**DataS1** Changes in protein abundance in roots and shoots of hydroponically grown Arabidopsis autophagy mutants compared to wild type.

**DataS2** Changes in protein abundance of proteins belonging to different subcellular locations in autophagy mutants compared to wild type.

**DataS3** Changes in protein degradation rate in autophagy mutants.

**DataS4** Changes in protein abundance and degradation rate in autophagy mutants.

**DataS5** Proteins with significant differences in protein abundance & degradation rate.

**DataS6** Root proteins with significant changes in their abundance & degradation rate.

**DataS7** Changes in protein abundance in wild type and autophagy mutants under control and Pi limiting conditions.

**DataS8** Changes in protein degradation rate in wild type and autophagy mutants under control and Pi limiting conditions.

**DataS9** Protein degradation rates and protein abundance data for PCA analysis.

**DataS10** Proteins with faster turnover rates under Pi limitation conditions in wild type

**DataS11** Metabolites measurement using mass spectrometry.

**DataS12** Precursor masses of metabolites used in LC-MS analysis.

## References

Ahn, H.K., Yoon, J.T., Choi, I., Kim, S., Lee, H.S., and Pai, H.S. (2019). Functional characterization of chaperonin containing T-complex polypeptide-1 and its conserved and novel substrates in Arabidopsis. Journal of experimental botany 70, 2741–2757.

Ames, B.N. (1966). [10] Assay of inorganic phosphate, total phosphate and phosphatases. In Methods in Enzymology (Academic Press), pp. 115–118.

An, H., and Harper, J.W. (2018). Systematic analysis of ribophagy in human cells reveals bystander flux during selective autophagy. Nat Cell Biol 20, 135–143.

Araújo, W.L., Tohge, T., Ishizaki, K., Leaver, C.J., and Fernie, A.R. (2011). Protein degradation – an alternative respiratory substrate for stressed plants. Trends Plant Sci.

Avin-Wittenberg, T., Bajdzienko, K., Wittenberg, G., Alseekh, S., Tohge, T., Bock, R., Giavalisco, P., and Fernie, A.R. (2015). Global analysis of the role of autophagy in cellular metabolism and energy homeostasis in Arabidopsis seedlings under carbon starvation. The Plant cell 27, 306–322.

Barros, J.A.S., Cavalcanti, J.H.F., Medeiros, D.B., Nunes-Nesi, A., Avin-Wittenberg, T., Fernie, A.R., and Araujo, W.L. (2017). Autophagy Deficiency Compromises Alternative Pathways of Respiration following Energy Deprivation in Arabidopsis thaliana. Plant physiology 175, 62–76.

Barros, J.A.S., Magen, S., Lapidot-Cohen, T., Rosental, L., Brotman, Y., Araujo, W.L., and Avin-Wittenberg, T. (2021). Autophagy is required for lipid homeostasis during dark-induced senescence. Plant physiology.

Bartsch, M., Bednarek, P., Vivancos, P.D., Schneider, B., von Roepenack-Lahaye, E., Foyer, C.H., Kombrink, E., Scheel, D., and Parker, J.E. (2010). Accumulation of Isochorismate-derived 2,3-Dihydroxybenzoic 3-O-beta-D-Xyloside in Arabidopsis Resistance to Pathogens and Ageing of Leaves. Journal of Biological Chemistry 285, 25654–25665.

Berson, T., von Wangenheim, D., Takac, T., Samajova, O., Rosero, A., Ovecka, M., Komis, G., Stelzer, E.H., and Samaj, J. (2014). Trans-Golgi network localized small GTPase RabA1d is involved in cell plate formation and oscillatory root hair growth. BMC Plant Biol 14, 252.

Bialecki, J.B., Ruzicka, J., Weisbecker, C.S., Haribal, M., and Attygalle, A.B. (2010). Collision-induced dissociation mass spectra of glucosinolate anions. J Mass Spectrom 45, 272–283.

Boyes, D.C., Zayed, A.M., Ascenzi, R., McCaskill, A.J., Hoffman, N.E., Davis, K.R., and Gorlach, J. (2001). Growth stage-based phenotypic analysis of Arabidopsis: a model for high throughput functional genomics in plants. The Plant cell 13, 1499–1510.

Chotewutmontri, P., and Barkan, A. (2016). Dynamics of Chloroplast Translation during Chloroplast Differentiation in Maize. PLoS Genet 12, e1006106.

Czechowski, T., Stitt, M., Altmann, T., Udvardi, M.K., and Scheible, W.R. (2005). Genome-wide identification and testing of superior reference genes for transcript normalization in Arabidopsis. Plant physiology 139, 5–17.

Dengjel, J., Høyer-Hansen, M., Nielsen, M.O., Eisenberg, T., Harder, L.M., Schandorff, S., Farkas, T., Kirkegaard, T., Becker, A.C., Schroeder, S., Vanselow, K., Lundberg, E., Nielsen, M.M., Kristensen, A.R., Akimov, V., Bunkenborg, J., Madeo, F., Jäättelä, M., and Andersen, J.S. (2012). Identification of Autophagosome-associated Proteins and Regulators by Quantitative Proteomic Analysis and Genetic Screens. Mol Cell Proteomics 11.

Deutsch, E.W., Mendoza, L., Shteynberg, D., Farrah, T., Lam, H., Tasman, N., Sun, Z., Nilsson, E., Pratt, B., Prazen, B., Eng, J.K., Martin, D.B., Nesvizhskii, A.I., and Aebersold, R. (2010). A guided tour of the Trans-Proteomic Pipeline. Proteomics 10, 1150–1159.

Farmer, L.M., Rinaldi, M.A., Young, P.G., Danan, C.H., Burkhart, S.E., and Bartel, B. (2013). Disrupting autophagy restores peroxisome function to an Arabidopsis lon2 mutant and reveals a role for the LON2 protease in peroxisomal matrix protein degradation. The Plant cell 25, 4085–4100.

Floyd, B.E., Morriss, S.C., MacIntosh, G.C., and Bassham, D.C. (2016). Evidence for autophagy-dependent pathways of rRNA turnover in Arabidopsis. Autophagy 11, 2199–2212.

Gao, C., Zhuang, X., Shen, J., and Jiang, L. (2017). Plant ESCRT Complexes: Moving Beyond Endosomal Sorting. Trends Plant Sci 22, 986–998.

Gao, J., Chaudhary, A., Vaddepalli, P., Nagel, M.K., Isono, E., and Schneitz, K. (2019). The Arabidopsis receptor kinase STRUBBELIG undergoes clathrin-dependent endocytosis. Journal of experimental botany 70, 3881–3894.

Gretzmeier, C., Eiselein, S., Johnson, G.R., Engelke, R., Nowag, H., Zarei, M., Kuttner, V., Becker, A.C., Rigbolt, K.T.G., Hoyer-Hansen, M., Andersen, J.S., Munz, C., Murphy, R.F., and Dengjel, J. (2017). Degradation of protein translation machinery by amino acid starvation-induced macroautophagy. Autophagy 13, 1064–1075.

Hamasaki, M., Noda, T., Baba, M., and Ohsumi, Y. (2005). Starvation triggers the delivery of the endoplasmic reticulum to the vacuole via autophagy in yeast. Traffic 6, 56–65.

Han, J., Gagnon, S., Eckle, T., and Borchers, C.H. (2013). Metabolomic analysis of key central carbon metabolism carboxylic acids as their 3-nitrophenylhydrazones by UPLC/ESI-MS. Electrophoresis 34, 2891–2900.

Han, S., Wang, Y., Zheng, X., Jia, Q., Zhao, J., Bai, F., Hong, Y., and Liu, Y. (2015). Cytoplastic Glyceraldehyde-3-Phosphate Dehydrogenases Interact with ATG3 to Negatively Regulate Autophagy and Immunity in Nicotiana benthamiana. The Plant cell 27, 1316–1331.

Have, M., Luo, J., Tellier, F., Balliau, T., Cueff, G., Chardon, F., Zivy, M., Rajjou, L., Cacas, J.L., and Masclaux-Daubresse, C. (2019). Proteomic and lipidomic analyses of the Arabidopsis atg5 autophagy mutant reveal major changes in endoplasmic reticulum and peroxisome metabolisms and in lipid composition. New Phytol 223, 1461–1477.

Henry, E., Fung, N., Liu, J., Drakakaki, G., and Coaker, G. (2015). Beyond glycolysis: GAPDHs are multi-functional enzymes involved in regulation of ROS, autophagy, and plant immune responses. PLoS Genet 11, e1005199.

Hohner, R., Marques, J.V., Ito, T., Amakura, Y., Budgeon, A.D., Jr., Weitz, K., Hixson, K.K., Davin, L.B., Kirchhoff, H., and Lewis, N.G. (2018). Reduced Arogenate Dehydratase Expression: Ramifications for Photosynthesis and Metabolism. Plant physiology 177, 115–131.

Hooper, C.M., Castleden, I.R., Tanz, S.K., Aryamanesh, N., and Millar, A.H. (2017). SUBA4: the interactive data analysis centre for Arabidopsis subcellular protein locations. Nucleic Acids Research 45, D1064–D1074.

Hooper, C.M., Tanz, S.K., Castleden, I.R., Vacher, M.A., Small, I.D., and Millar, A.H. (2014). SUBAcon: a consensus algorithm for unifying the subcellular localization data of the Arabidopsis proteome. Bioinformatics 30, 3356–3364.

Huang, X., Zheng, C., Liu, F., Yang, C., Zheng, P., Lu, X., Tian, J., Chung, T., Otegui, M.S., Xiao, S., Gao, C., Vierstra, R.D., and Li, F. (2019). Genetic Analyses of the Arabidopsis ATG1 Kinase Complex Reveal Both Kinase-Dependent and Independent Autophagic Routes during Fixed-Carbon Starvation. The Plant cell 31, 2973–2995.

Izumi, M., Ishida, H., Nakamura, S., and Hidema, J. (2017). Entire Photodamaged Chloroplasts Are Transported to the Central Vacuole by Autophagy. The Plant cell 29, 377–394.

Jiao, L., Zhang, H.L., Li, D.D., Yang, K.L., Tang, J., Li, X., Ji, J., Yu, Y., Wu, R.Y., Ravichandran, S., Liu, J.J., Feng, G.K., Chen, M.S., Zeng, Y.X., Deng, R., and Zhu, X.F. (2017). Regulation of Glycolytic Metabolism by Autophagy in Liver Cancer Involves Selective Autophagic Degradation of HK2 (hexokinase 2). Autophagy, 0.

Jung, H., Lee, H.N., Marshall, R.S., Lomax, A.W., Yoon, M.J., Kim, J., Kim, J.H., Vierstra, R.D., Chung, T., and Bozhkov, P. (2020). Arabidopsis cargo receptor NBR1 mediates selective autophagy of defective proteins. Journal of experimental botany 71, 73–89.

Juntawong, P., Girke, T., Bazin, J., and Bailey-Serres, J. (2014). Translational dynamics revealed by genome-wide profiling of ribosome footprints in Arabidopsis. Proceedings of the National Academy of Sciences of the United States of America 111, E203–212.

Kast, D.J., and Dominguez, R. (2017). The Cytoskeleton–Autophagy Connection. Current Biology 27, R318–R326.

Khaminets, A., Heinrich, T., Mari, M., Grumati, P., Huebner, A.K., Akutsu, M., Liebmann, L., Stolz, A., Nietzsche, S., Koch, N., Mauthe, M., Katona, I., Qualmann, B., Weis, J., Reggiori, F., Kurth, I., Hubner, C.A., and Dikic, I. (2015). Regulation of endoplasmic reticulum turnover by selective autophagy. Nature 522, 354–358.

Lam, S.K., Cai, Y., Tse, Y.C., Wang, J., Law, A.H., Pimpl, P., Chan, H.Y., Xia, J., and Jiang, L. (2009). BFA-induced compartments from the Golgi apparatus and trans-Golgi network/early endosome are distinct in plant cells. Plant J 60, 865–881.

Lamesch, P., Berardini, T.Z., Li, D., Swarbreck, D., Wilks, C., Sasidharan, R., Muller, R., Dreher, K., Alexander, D.L., Garcia-Hernandez, M., Karthikeyan, A.S., Lee, C.H., Nelson, W.D., Ploetz, L., Singh, S., Wensel, A., and Huala, E. (2012). The Arabidopsis Information Resource (TAIR): improved gene annotation and new tools. Nucleic Acids Res 40, D1202–1210.

Lee, K.C., Chan, W., Liang, Z.T., Liu, N., Zhao, Z.Z., Lee, A.W.M., and Cai, Z.W. (2008). Rapid screening method for intact glucosinolates in Chinese medicinal herbs by using liquid chromatography coupled with electrospray ionization ion trap mass spectrometry in negative ion mode. Rapid Commun Mass Sp 22, 2825–2834.

Li, F.Q., and Vierstra, R.D. (2014). Arabidopsis ATG11, a scaffold that links the ATG1-ATG13 kinase complex to general autophagy and selective mitophagy. Autophagy 10, 1466–1467.

Li, F.Q., Chung, T., and Vierstra, R.D. (2014). AUTOPHAGY-RELATED11 Plays a Critical Role in General Autophagy- and Senescence-Induced Mitophagy in Arabidopsis. The Plant cell 26, 788–807.

Li, L., Nelson, C.J., Trosch, J., Castleden, I., Huang, S., and Millar, A.H. (2017). Protein Degradation Rate in Arabidopsis thaliana Leaf Growth and Development. Plant Cell 29, 207–228.

Lin, L.Z., Sun, J.H., Chen, P., Zhang, R.W., Fan, X.E., Li, L.W., and Harnly, J.M. (2014). Profiling of Glucosinolates and Flavonoids in Rorippa indica (Linn.) Hiern. (Cruciferae) by UHPLC-PDA-ESI/HRMSn. J Agr Food Chem 62, 6118–6129.

Liu, L., Feng, D., Chen, G., Chen, M., Zheng, Q., Song, P., Ma, Q., Zhu, C., Wang, R., Qi, W., Huang, L., Xue, P., Li, B., Wang, X., Jin, H., Wang, J., Yang, F., Liu, P., Zhu, Y., Sui, S., and Chen, Q. (2012a). Mitochondrial outer-membrane protein FUNDC1 mediates hypoxia-induced mitophagy in mammalian cells. Nat Cell Biol 14, 177–185.

Liu, Y., Burgos, J.S., Deng, Y., Srivastava, R., Howell, S.H., and Bassham, D.C. (2012b). Degradation of the endoplasmic reticulum by autophagy during endoplasmic reticulum stress in Arabidopsis. The Plant cell 24, 4635–4651.

Macharia, M.W., Tan, W.Y.Z., Das, P.P., Naqvi, N.I., and Wong, S.M. (2019). Proximity-dependent biotinylation screening identifies NbHYPK as a novel interacting partner of ATG8 in plants. BMC Plant Biol 19, 326.

Mackeh, R., Perdiz, D., Lorin, S., Codogno, P., and Pous, C. (2013). Autophagy and microtubules - new story, old players. J Cell Sci 126, 1071–1080.

Marshall, R.S., and Vierstra, R.D. (2018). Autophagy: The Master of Bulk and Selective Recycling. Annu Rev Plant Biol 69, 173–208.

Marshall, R.S., Li, F., Gemperline, D.C., Book, A.J., and Vierstra, R.D. (2015). Autophagic Degradation of the 26S Proteasome Is Mediated by the Dual ATG8/Ubiquitin Receptor RPN10 in Arabidopsis. Molecular cell 58, 1053–1066.

Marty, F. (1999). Plant vacuoles. The Plant cell 11, 587–600.

Masclaux-Daubresse, C., Clement, G., Anne, P., Routaboul, J.M., Guiboileau, A., Soulay, F., Shirasu, K., and Yoshimoto, K. (2014). Stitching together the Multiple Dimensions of Autophagy Using Metabolomics and Transcriptomics Reveals Impacts on Metabolism, Development, and Plant Responses to the Environment in Arabidopsis. The Plant cell 26, 1857–1877.

Matsuda, F., Yonekura-Sakakibara, K., Niida, R., Kuromori, T., Shinozaki, K., and Saito, K. (2009). MS/MS spectral tag-based annotation of non-targeted profile of plant secondary metabolites. Plant Journal 57, 555–577.

McLoughlin, F., Augustine, R.C., Marshall, R.S., Li, F., Kirkpatrick, L.D., Otegui, M.S., and Vierstra, R.D. (2018). Maize multi-omics reveal roles for autophagic recycling in proteome remodelling and lipid turnover. Nat Plants.

McLoughlin, F., Marshall, R.S., Ding, X., Chatt, E.C., Kirkpatrick, L.D., Augustine, R.C., Li, F., Otegui, M.S., and Vierstra, R.D. (2020). Autophagy Plays Prominent Roles in Amino Acid, Nucleotide, and Carbohydrate Metabolism during Fixed-Carbon Starvation in Maize. The Plant cell 32, 2699–2724.

McWhite, C.D., Papoulas, O., Drew, K., Cox, R.M., June, V., Dong, O.X., Kwon, T., Wan, C., Salmi, M.L., Roux, S.J., Browning, K.S., Chen, Z.J., Ronald, P.C., and Marcotte, E.M. (2020). A Pan-plant Protein Complex Map Reveals Deep Conservation and Novel Assemblies. Cell 181, 460–474.e414.

Monastyrska, I., Rieter, E., Klionsky, D.J., and Reggiori, F. (2009). Multiple roles of the cytoskeleton in autophagy. Biol Rev Camb Philos Soc 84, 431–448.

Naumann, C., Muller, J., Sakhonwasee, S., Wieghaus, A., Hause, G., Heisters, M., Burstenbinder, K., and Abel, S. (2019). The Local Phosphate Deficiency Response Activates Endoplasmic Reticulum Stress-Dependent Autophagy. Plant physiology 179, 460–476.

Nebenfuhr, A., Ritzenthaler, C., and Robinson, D.G. (2002). Brefeldin A: deciphering an enigmatic inhibitor of secretion. Plant physiology 130, 1102–1108.

Nelson, C.J., Alexova, R., Jacoby, R.P., and Millar, A.H. (2014). Proteins with high turnover rate in barley leaves estimated by proteome analysis combined with in planta isotope labeling. Plant physiology 166, 91–108.

Nesvizhskii, A.I., Keller, A., Kolker, E., and Aebersold, R. (2003). A statistical model for identifying proteins by tandem mass spectrometry. Anal Chem 75, 4646–4658.

Otegui, M.S. (2018). Vacuolar degradation of chloroplast components: autophagy and beyond. Journal of experimental botany 69, 741–750.

Pavel, M., Imarisio, S., Menzies, F.M., Jimenez-Sanchez, M., Siddiqi, F.H., Wu, X., Renna, M., O’Kane, C.J., Crowther, D.C., and Rubinsztein, D.C. (2016). CCT complex restricts neuropathogenic protein aggregation via autophagy. Nature communications 7, 13821.

Pereira, C., Pereira, S., and Pissarra, J. (2014). Delivering of proteins to the plant vacuole--an update. Int J Mol Sci 15, 7611–7623.

Qian, X., Li, X., and Lu, Z. (2017a). Protein kinase activity of the glycolytic enzyme PGK1 regulates autophagy to promote tumorigenesis. Autophagy 13, 1246–1247.

Qian, X., Li, X.J., Cai, Q.S., Zhang, C.B., Yu, Q.J., Jiang, Y.H., Lee, J.H., Hawke, D., Wang, Y.G., Xia, Y., Zheng, Y., Jiang, B.H., Liu, D.X., Jiang, T., and Lu, Z.M. (2017b). Phosphoglycerate Kinase 1 Phosphorylates Beclin1 to Induce Autophagy. Molecular cell 65, 917–+.

Razi, M., Chan, E.Y., and Tooze, S.A. (2009). Early endosomes and endosomal coatomer are required for autophagy. J Cell Biol 185, 305–321.

Reggiori, F., Monastyrska, I., Shintani, T., and Klionsky, D.J. (2005). The actin cytoskeleton is required for selective types of autophagy, but not nonspecific autophagy, in the yeast Saccharomyces cerevisiae. Mol Biol Cell 16, 5843–5856.

Rochfort, S.J., Trenerry, V.C., Imsic, M., Panozzo, J., and Jones, R. (2008). Class targeted metabolomics: ESI ion trap screening methods for glucosinolates based on MSn fragmentation. Phytochemistry 69, 1671–1679.

Schindelin, J., Arganda-Carreras, I., Frise, E., Kaynig, V., Longair, M., Pietzsch, T., Preibisch, S., Rueden, C., Saalfeld, S., Schmid, B., Tinevez, J.Y., White, D.J., Hartenstein, V., Eliceiri, K., Tomancak, P., and Cardona, A. (2012). Fiji: an open-source platform for biological-image analysis. Nat Methods 9, 676–682.

Shimada, T., Takagi, J., Ichino, T., Shirakawa, M., and Hara-Nishimura, I. (2018). Plant Vacuoles. Annu Rev Plant Biol 69, 123–145.

Shteynberg, D., Deutsch, E.W., Lam, H., Eng, J.K., Sun, Z., Tasman, N., Mendoza, L., Moritz, R.L., Aebersold, R., and Nesvizhskii, A.I. (2011). iProphet: multi-level integrative analysis of shotgun proteomic data improves peptide and protein identification rates and error estimates. Molecular & cellular proteomics : MCP 10, M111 007690.

Stobiecki, M., Skirycz, A., Kerhoas, L., Kachlicki, P., Muth, D., Einhorn, J., and Mueller-Roeber, B. (2006). Profiling of phenolic glycosidic conjugates in leaves of Arabidopsis thaliana using LC/MS. Metabolomics 2, 197–219.

Styers, M.L., O’Connor, A.K., Grabski, R., Cormet-Boyaka, E., and Sztul, E. (2008). Depletion of beta-COP reveals a role for COP-I in compartmentalization of secretory compartments and in biosynthetic transport of caveolin-1. Am J Physiol Cell Physiol 294, C1485–1498.

Sugiyama, R., and Hirai, M.Y. (2019). Atypical Myrosinase as a Mediator of Glucosinolate Functions in Plants. Frontiers in plant science 10.

Tasaki, M., Asatsuma, S., and Matsuoka, K. (2014). Monitoring protein turnover during phosphate starvation-dependent autophagic degradation using a photoconvertible fluorescent protein aggregate in tobacco BY-2 cells. Frontiers in plant science 5, 172.

Thompson, A.R., Doelling, J.H., Suttangkakul, A., and Vierstra, R.D. (2005). Autophagic nutrient recycling in Arabidopsis directed by the ATG8 and ATG12 conjugation pathways. Plant physiology 138, 2097–2110.

Thormahlen, I., Zupok, A., Rescher, J., Leger, J., Weissenberger, S., Groysman, J., Orwat, A., Chatel-Innocenti, G., Issakidis-Bourguet, E., Armbruster, U., and Geigenberger, P. (2017). Thioredoxins Play a Crucial Role in Dynamic Acclimation of Photosynthesis in Fluctuating Light. Mol Plant 10, 168–182.

Vandesompele, J., De Preter, K., Pattyn, F., Poppe, B., Van Roy, N., De Paepe, A., and Speleman, F. (2002). Accurate normalization of real-time quantitative RT-PCR data by geometric averaging of multiple internal control genes. Genome Biol 3, RESEARCH0034.

Vincow, E.S., Thomas, R.E., Merrihew, G.E., Shulman, N.J., Bammler, T.K., MacDonald, J.W., MacCoss, M.J., and Pallanck, L.J. (2019). Autophagy accounts for approximately one-third of mitochondrial protein turnover and is protein selective. Autophagy 15, 1592–1605.

Wang, J., Zhang, J., Lee, Y.M., Koh, P.L., Ng, S., Bao, F., Lin, Q., and Shen, H.M. (2016). Quantitative chemical proteomics profiling of de novo protein synthesis during starvation-mediated autophagy. Autophagy 12, 1931–1944.

Wang, S., Xie, K., Xu, G., Zhou, H., Guo, Q., Wu, J., Liao, Z., Liu, N., Wang, Y., and Liu, Y. (2018). Plant G proteins interact with endoplasmic reticulum luminal protein receptors to regulate endoplasmic reticulum retrieval. J Integr Plant Biol 60, 541–561.

Waters, M.T., Nelson, D.C., Scaffidi, A., Flematti, G.R., Sun, Y.K., Dixon, K.W., and Smith, S.M. (2012). Specialisation within the DWARF14 protein family confers distinct responses to karrikins and strigolactones in Arabidopsis. Development 139, 1285–1295.

Watson, A.S., Riffelmacher, T., Stranks, A., Williams, O., De Boer, J., Cain, K., MacFarlane, M., McGouran, J., Kessler, B., Khandwala, S., Chowdhury, O., Puleston, D., Phadwal, K., Mortensen, M., Ferguson, D., Soilleux, E., Woll, P., Jacobsen, S.E., and Simon, A.K. (2015). Autophagy limits proliferation and glycolytic metabolism in acute myeloid leukemia. Cell Death Discov 1.

Wijerathna-Yapa, A., Stroeher, E., Fenske, R., Li, L., Duncan, O., and Millar, A.H. (2021). Proteomics for Autophagy Receptor and Cargo Identification in Plants. J Proteome Res 20, 129–138.

Xiong, Y., Contento, A.L., and Bassham, D.C. (2005). AtATG18a is required for the formation of autophagosomes during nutrient stress and senescence in Arabidopsis thaliana. Plant J 42, 535–546.

Xu, J., Kozlov, G., McPherson, P.S., and Gehring, K. (2018). A PH-like domain of the Rab12 guanine nucleotide exchange factor DENND3 binds actin and is required for autophagy. J Biol Chem 293, 4566–4574.

Yokota, H., Gomi, K., and Shintani, T. (2017). Induction of autophagy by phosphate starvation in an Atg11-dependent manner in Saccharomyces cerevisiae. Biochem Biophys Res Commun 483, 522–527.

Yoshimoto, K., Ishida, H., Wada, S., Ohsumi, Y., and Shirasu, K. (2009a). The role of plant autophagy in nutrient starvation and aging. Autophagy 5, 904–904.

Yoshimoto, K., Jikumaru, Y., Kamiya, Y., Kusano, M., Consonni, C., Panstruga, R., Ohsumi, Y., and Shirasu, K. (2009b). Autophagy negatively regulates cell death by controlling NPR1-dependent salicylic acid signaling during senescence and the innate immune response in Arabidopsis. The Plant cell 21, 2914–2927.

Zhang, C., Hicks, G.R., and Raikhel, N.V. (2014). Plant vacuole morphology and vacuolar trafficking. Frontiers in plant science 5.

Zhang, K., Halitschke, R., Yin, C., Liu, C.J., and Gan, S.S. (2013). Salicylic acid 3-hydroxylase regulates Arabidopsis leaf longevity by mediating salicylic acid catabolism. Proc Natl Acad Sci U S A 110, 14807–14812.

Zhang, T., Shen, S., Qu, J., and Ghaemmaghami, S. (2016). Global Analysis of Cellular Protein Flux Quantifies the Selectivity of Basal Autophagy. Cell Reports 14, 2426–2439.

Zhang, X., Ding, X., Marshall, R.S., Paez-Valencia, J., Lacey, P., Vierstra, R.D., and Otegui, M.S. (2020). Reticulon proteins modulate autophagy of the endoplasmic reticulum in maize endosperm. Elife 9.

Zheng, X., Wu, M., Li, X., Cao, J., Li, J., Wang, J., Huang, S., Liu, Y., and Wang, Y. (2019). Actin filaments are dispensable for bulk autophagy in plants. Autophagy 15, 2126–2141.

Zhuang, X., and Jiang, L. (2019). Chloroplast Degradation: Multiple Routes Into the Vacuole. Frontiers in plant science 10, 359.

## Parsed Citations

Ahn, H.K., Yoon, J.T., Choi, I., Kim, S., Lee, H.S., and Pai, H.S. (2019). Functional characterization of chaperonin containing T-complex polypeptide-1 and its conserved and novel substrates in Arabidopsis. Journal of experimental botany 70, 2741–2757. Google Scholar: Author Only Title Only Author and Title

Ames, B.N. (1966). [10] Assay of inorganic phosphate, total phosphate and phosphatases. In Methods in Enzymology (Academic Press), pp. 115–118. Google Scholar: Author Only Title Only Author and Title

An, H., and Harper, J.W. (2018). Systematic analysis of ribophagy in human cells reveals bystander flux during selective autophagy. Nat Cell Biol 20, 135–143. Google Scholar: Author Only Title Only Author and Title

Araújo, W.L., Tohge, T., Ishizaki, K., Leaver, C.J., and Fernie, A.R. (2011). Protein degradation – an alternative respiratory substrate for stressed plants. Trends Plant Sci.Google Scholar: Author Only Title Only Author and Title

Avin-Wittenberg, T., Bajdzienko, K., Wittenberg, G., Alseekh, S., Tohge, T., Bock, R., Giavalisco, P., and Fernie, A.R. (2015). Global analysis of the role of autophagy in cellular metabolism and energy homeostasis in Arabidopsis seedlings under carbon starvation. The Plant cell 27, 306–322. Google Scholar: Author Only Title Only Author and Title

Barros, J.A.S., Cavalcanti, J.H.F., Medeiros, D.B., Nunes-Nesi, A., Avin-Wittenberg, T., Fernie, A.R., and Araujo, W.L. (2017). Autophagy Deficiency Compromises Alternative Pathways of Respiration following Energy Deprivation in Arabidopsis thaliana. Plant physiology 175, 62–76. Google Scholar: Author Only Title Only Author and Title

Barros, J.A.S., Magen, S., Lapidot-Cohen, T., Rosental, L., Brotman, Y., Araujo, W.L., and Avin-Wittenberg, T. (2021). Autophagy is required for lipid homeostasis during dark-induced senescence. Plant physiology. Google Scholar: Author Only Title Only Author and Title

Bartsch, M., Bednarek, P., Vivancos, P.D., Schneider, B., von Roepenack-Lahaye, E., Foyer, C.H., Kombrink, E., Scheel, D., and Parker, J.E. (2010). Accumulation of Isochorismate-derived 2,3-Dihydroxybenzoic 3-O-beta-D-Xyloside in Arabidopsis Resistance to Pathogens and Ageing of Leaves. Journal of Biological Chemistry 285, 25654–25665. Google Scholar: Author Only Title Only Author and Title

Berson, T., von Wangenheim, D., Takac, T., Samajova, O., Rosero, A., Ovecka, M., Komis, G., Stelzer, E.H., and Samaj, J. (2014). Trans-Golgi network localized small GTPase RabA1d is involved in cell plate formation and oscillatory root hair growth. BMC Plant Biol 14, 252. Google Scholar: Author Only Title Only Author and Title

Bialecki, J.B., Ruzicka, J., Weisbecker, C.S., Haribal, M., and Attygalle, A.B. (2010). Collision-induced dissociation mass spectra of glucosinolate anions. J Mass Spectrom 45, 272–283. Google Scholar: Author Only Title Only Author and Title

Boyes, D.C., Zayed, A.M., Ascenzi, R., McCaskill, A.J., Hoffman, N.E., Davis, K.R., and Gorlach, J. (2001). Growth stage-based phenotypic analysis of Arabidopsis: a model for high throughput functional genomics in plants. The Plant cell 13, 1499–1510. Google Scholar: Author Only Title Only Author and Title

Chotewutmontri, P., and Barkan, A. (2016). Dynamics of Chloroplast Translation during Chloroplast Differentiation in Maize. PLoS Genet 12, e1006106. Google Scholar: Author Only Title Only Author and Title

Czechowski, T., Stitt, M., Altmann, T., Udvardi, M.K., and Scheible, W.R. (2005). Genome-wide identification and testing of superior reference genes for transcript normalization in Arabidopsis. Plant physiology 139, 5–17. Google Scholar: Author Only Title Only Author and Title

Dengjel, J., Høyer-Hansen, M., Nielsen, M.O., Eisenberg, T., Harder, L.M., Schandorff, S., Farkas, T., Kirkegaard, T., Becker, A.C., Schroeder, S., Vanselow, K., Lundberg, E., Nielsen, M.M., Kristensen, A.R., Akimov, V., Bunkenborg, J., Madeo, F., Jäättelä, M., and Andersen, J.S. (2012). Identification of Autophagosome-associated Proteins and Regulators by Quantitative Proteomic Analysis and Genetic Screens. Mol Cell Proteomics 11. Google Scholar: Author Only Title Only Author and Title

Deutsch, E.W., Mendoza, L., Shteynberg, D., Farrah, T., Lam, H., Tasman, N., Sun, Z., Nilsson, E., Pratt, B., Prazen, B., Eng, J.K., Martin, D.B., Nesvizhskii, A.I., and Aebersold, R. (2010). A guided tour of the Trans-Proteomic Pipeline. Proteomics 10, 1150–1159. Google Scholar: Author Only Title Only Author and Title

Farmer, L.M., Rinaldi, M.A., Young, P.G., Danan, C.H., Burkhart, S.E., and Bartel, B. (2013). Disrupting autophagy restores peroxisome function to an Arabidopsis lon2 mutant and reveals a role for the LON2 protease in peroxisomal matrix protein degradation. The Plant cell 25, 4085–4100. Google Scholar: Author Only Title Only Author and Title

Floyd, B.E., Morriss, S.C., MacIntosh, G.C., and Bassham, D.C. (2016). Evidence for autophagy-dependent pathways of rRNA turnover in Arabidopsis. Autophagy 11, 2199–2212. Google Scholar: Author Only Title Only Author and Title

Gao, C., Zhuang, X., Shen, J., and Jiang, L. (2017). Plant ESCRT Complexes: Moving Beyond Endosomal Sorting. Trends Plant Sci 22, 986–998. Google Scholar: Author Only Title Only Author and Title

Gao, J., Chaudhary, A., Vaddepalli, P., Nagel, M.K., Isono, E., and Schneitz, K. (2019). The Arabidopsis receptor kinase STRUBBELIG undergoes clathrin-dependent endocytosis. Journal of experimental botany 70, 3881–3894. Google Scholar: Author Only Title Only Author and Title

Gretzmeier, C., Eiselein, S., Johnson, G.R., Engelke, R., Nowag, H., Zarei, M., Kuttner, V., Becker, A.C., Rigbolt, K.T.G., Hoyer-Hansen, M., Andersen, J.S., Munz, C., Murphy, R.F., and Dengjel, J. (2017). Degradation of protein translation machinery by amino acid starvation-induced macroautophagy. Autophagy 13, 1064–1075. Google Scholar: Author Only Title Only Author and Title

Hamasaki, M., Noda, T., Baba, M., and Ohsumi, Y. (2005). Starvation triggers the delivery of the endoplasmic reticulum to the vacuole via autophagy in yeast. Traffic 6, 56–65. Google Scholar: Author Only Title Only Author and Title

Han, J., Gagnon, S., Eckle, T., and Borchers, C.H. (2013). Metabolomic analysis of key central carbon metabolism carboxylic acids as their 3-nitrophenylhydrazones by UPLC/ESI-MS. Electrophoresis 34, 2891–2900. Google Scholar: Author Only Title Only Author and Title

Han, S., Wang, Y., Zheng, X., Jia, Q., Zhao, J., Bai, F., Hong, Y., and Liu, Y. (2015). Cytoplastic Glyceraldehyde-3-Phosphate Dehydrogenases Interact with ATG3 to Negatively Regulate Autophagy and Immunity in Nicotiana benthamiana. The Plant cell 27, 1316–1331. Google Scholar: Author Only Title Only Author and Title

Have, M., Luo, J., Tellier, F., Balliau, T., Cueff, G., Chardon, F., Zivy, M., Rajjou, L., Cacas, J.L., and Masclaux-Daubresse, C. (2019). Proteomic and lipidomic analyses of the Arabidopsis atg5 autophagy mutant reveal major changes in endoplasmic reticulum and peroxisome metabolisms and in lipid composition. New Phytol 223, 1461–1477. Google Scholar: Author Only Title Only Author and Title

Henry, E., Fung, N., Liu, J., Drakakaki, G., and Coaker, G. (2015). Beyond glycolysis: GAPDHs are multi-functional enzymes involved in regulation of ROS, autophagy, and plant immune responses. PLoS Genet 11, e1005199. Google Scholar: Author Only Title Only Author and Title

Hohner, R., Marques, J.V., Ito, T., Amakura, Y., Budgeon, A.D., Jr., Weitz, K., Hixson, K.K., Davin, L.B., Kirchhoff, H., and Lewis, N.G. (2018). Reduced Arogenate Dehydratase Expression: Ramifications for Photosynthesis and Metabolism. Plant physiology 177, 115–131. Google Scholar: Author Only Title Only Author and Title

Hooper, C.M., Castleden, I.R., Tanz, S.K., Aryamanesh, N., and Millar, A.H. (2017). SUBA4: the interactive data analysis centre for Arabidopsis subcellular protein locations. Nucleic Acids Research 45, D1064–D1074. Google Scholar: Author Only Title Only Author and Title

Hooper, C.M., Tanz, S.K., Castleden, I.R., Vacher, M.A., Small, I.D., and Millar, A.H. (2014). SUBAcon: a consensus algorithm for unifying the subcellular localization data of the Arabidopsis proteome. Bioinformatics 30, 3356–3364. Google Scholar: Author Only Title Only Author and Title

Huang, X., Zheng, C., Liu, F., Yang, C., Zheng, P., Lu, X., Tian, J., Chung, T., Otegui, M.S., Xiao, S., Gao, C., Vierstra, R.D., and Li, F. (2019). Genetic Analyses of the Arabidopsis ATG1 Kinase Complex Reveal Both Kinase-Dependent and Independent Autophagic Routes during Fixed-Carbon Starvation. The Plant cell 31, 2973–2995. Google Scholar: Author Only Title Only Author and Title

Izumi, M., Ishida, H., Nakamura, S., and Hidema, J. (2017). Entire Photodamaged Chloroplasts Are Transported to the Central Vacuole by Autophagy. The Plant cell 29, 377–394. Google Scholar: Author Only Title Only Author and Title

Jiao, L., Zhang, H.L., Li, D.D., Yang, K.L., Tang, J., Li, X., Ji, J., Yu, Y., Wu, R.Y., Ravichandran, S., Liu, J.J., Feng, G.K., Chen, M.S., Zeng, Y.X., Deng, R., and Zhu, X.F. (2017). Regulation of Glycolytic Metabolism by Autophagy in Liver Cancer Involves Selective Autophagic Degradation of HK2 (hexokinase 2). Autophagy, 0. Google Scholar: Author Only Title Only Author and Title

Jung, H., Lee, H.N., Marshall, R.S., Lomax, A.W., Yoon, M.J., Kim, J., Kim, J.H., Vierstra, R.D., Chung, T., and Bozhkov, P. (2020). Arabidopsis cargo receptor NBR1 mediates selective autophagy of defective proteins. Journal of experimental botany 71, 73–89. Google Scholar: Author Only Title Only Author and Title

Juntawong, P., Girke, T., Bazin, J., and Bailey-Serres, J. (2014). Translational dynamics revealed by genome-wide profiling of ribosome footprints in Arabidopsis. Proceedings of the National Academy of Sciences of the United States of America 111, E203–212. Google Scholar: Author Only Title Only Author and Title

Kast, D.J., and Dominguez, R. (2017). The Cytoskeleton–Autophagy Connection. Current Biology 27, R318–R326. Google Scholar: Author Only Title Only Author and Title

Khaminets, A., Heinrich, T., Mari, M., Grumati, P., Huebner, A.K., Akutsu, M., Liebmann, L., Stolz, A., Nietzsche, S., Koch, N., Mauthe, M., Katona, I., Qualmann, B., Weis, J., Reggiori, F., Kurth, I., Hubner, C.A., and Dikic, I. (2015). Regulation of endoplasmic reticulum turnover by selective autophagy. Nature 522, 354–358. Google Scholar: Author Only Title Only Author and Title

Lam, S.K., Cai, Y., Tse, Y.C., Wang, J., Law, A.H., Pimpl, P., Chan, H.Y., Xia, J., and Jiang, L. (2009). BFA-induced compartments from the Golgi apparatus and trans-Golgi network/early endosome are distinct in plant cells. Plant J 60, 865–881. Google Scholar: Author Only Title Only Author and Title

Lamesch, P., Berardini, T.Z., Li, D., Swarbreck, D., Wilks, C., Sasidharan, R., Muller, R., Dreher, K., Alexander, D.L., Garcia-Hernandez, M., Karthikeyan, A.S., Lee, C.H., Nelson, W.D., Ploetz, L., Singh, S., Wensel, A., and Huala, E. (2012). The Arabidopsis Information Resource (TAIR): improved gene annotation and new tools. Nucleic Acids Res 40, D1202–1210. Google Scholar: Author Only Title Only Author and Title

Lee, K.C., Chan, W., Liang, Z.T., Liu, N., Zhao, Z.Z., Lee, A.W.M., and Cai, Z.W. (2008). Rapid screening method for intact glucosinolates in Chinese medicinal herbs by using liquid chromatography coupled with electrospray ionization ion trap mass spectrometry in negative ion mode. Rapid Commun Mass Sp 22, 2825–2834. Google Scholar: Author Only Title Only Author and Title

Li, F.Q., and Vierstra, R.D. (2014). Arabidopsis ATG11, a scaffold that links the ATG1-ATG13 kinase complex to general autophagy and selective mitophagy. Autophagy 10, 1466–1467. Google Scholar: Author Only Title Only Author and Title

Li, F.Q., Chung, T., and Vierstra, R.D. (2014). AUTOPHAGY-RELATED11 Plays a Critical Role in General Autophagy- and Senescence-Induced Mitophagy in Arabidopsis. The Plant cell 26, 788–807. Google Scholar: Author Only Title Only Author and Title

Li, L., Nelson, C.J., Trosch, J., Castleden, I., Huang, S., and Millar, A.H. (2017). Protein Degradation Rate in Arabidopsis thaliana Leaf Growth and Development. Plant Cell 29, 207–228. Google Scholar: Author Only Title Only Author and Title

Lin, L.Z., Sun, J.H., Chen, P., Zhang, R.W., Fan, X.E., Li, L.W., and Harnly, J.M. (2014). Profiling of Glucosinolates and Flavonoids in Rorippa indica (Linn.) Hiern. (Cruciferae) by UHPLC-PDA-ESI/HRMSn. J Agr Food Chem 62, 6118–6129. Google Scholar: Author Only Title Only Author and Title

Liu, L., Feng, D., Chen, G., Chen, M., Zheng, Q., Song, P., Ma, Q., Zhu, C., Wang, R., Qi, W., Huang, L., Xue, P., Li, B., Wang, X., Jin, H., Wang, J., Yang, F., Liu, P., Zhu, Y., Sui, S., and Chen, Q. (2012a). Mitochondrial outer-membrane protein FUNDC1 mediates hypoxia-induced mitophagy in mammalian cells. Nat Cell Biol 14, 177–185. Google Scholar: Author Only Title Only Author and Title

Liu, Y., Burgos, J.S., Deng, Y., Srivastava, R., Howell, S.H., and Bassham, D.C. (2012b). Degradation of the endoplasmic reticulum by autophagy during endoplasmic reticulum stress in Arabidopsis. The Plant cell 24, 4635–4651. Google Scholar: Author Only Title Only Author and Title

Macharia, M.W., Tan, W.Y.Z., Das, P.P., Naqvi, N.I., and Wong, S.M. (2019). Proximity-dependent biotinylation screening identifies NbHYPK as a novel interacting partner of ATG8 in plants. BMC Plant Biol 19, 326. Google Scholar: Author Only Title Only Author and Title

Mackeh, R., Perdiz, D., Lorin, S., Codogno, P., and Pous, C. (2013). Autophagy and microtubules – new story, old players. J Cell Sci 126, 1071–1080. Google Scholar: Author Only Title Only Author and Title

Marshall, R.S., and Vierstra, R.D. (2018). Autophagy: The Master of Bulk and Selective Recycling. Annu Rev Plant Biol 69, 173–208. Google Scholar: Author Only Title Only Author and Title

Marshall, R.S., Li, F., Gemperline, D.C., Book, A.J., and Vierstra, R.D. (2015). Autophagic Degradation of the 26S Proteasome Is Mediated by the Dual ATG8/Ubiquitin Receptor RPN10 in Arabidopsis. Molecular cell 58, 1053–1066. Google Scholar: Author Only Title Only Author and Title

Marty, F. (1999). Plant vacuoles. The Plant cell 11, 587–600. Google Scholar: Author Only Title Only Author and Title

Masclaux-Daubresse, C., Clement, G., Anne, P., Routaboul, J.M., Guiboileau, A., Soulay, F., Shirasu, K., and Yoshimoto, K. (2014). Stitching together the Multiple Dimensions of Autophagy Using Metabolomics and Transcriptomics Reveals Impacts on Metabolism, Development, and Plant Responses to the Environment in Arabidopsis. The Plant cell 26, 1857–1877. Google Scholar: Author Only Title Only Author and Title

Matsuda, F., Yonekura-Sakakibara, K., Niida, R., Kuromori, T., Shinozaki, K., and Saito, K. (2009). MS/MS spectral tag-based annotation of non-targeted profile of plant secondary metabolites. Plant Journal 57, 555–577. Google Scholar: Author Only Title Only Author and Title

McLoughlin, F., Augustine, R.C., Marshall, R.S., Li, F., Kirkpatrick, L.D., Otegui, M.S., and Vierstra, R.D. (2018). Maize multi-omics reveal roles for autophagic recycling in proteome remodelling and lipid turnover. Nat Plants. Google Scholar: Author Only Title Only Author and Title

McLoughlin, F., Marshall, R.S., Ding, X., Chatt, E.C., Kirkpatrick, L.D., Augustine, R.C., Li, F., Otegui, M.S., and Vierstra, R.D. (2020). Autophagy Plays Prominent Roles in Amino Acid, Nucleotide, and Carbohydrate Metabolism during Fixed-Carbon Starvation in Maize. The Plant cell 32, 2699–2724. Google Scholar: Author Only Title Only Author and Title

McWhite, C.D., Papoulas, O., Drew, K., Cox, R.M., June, V., Dong, O.X., Kwon, T., Wan, C., Salmi, M.L., Roux, S.J., Browning, K.S., Chen, Z.J., Ronald, P.C., and Marcotte, E.M. (2020). A Pan-plant Protein Complex Map Reveals Deep Conservation and Novel Assemblies. Cell 181, 460–474.e414. Google Scholar: Author Only Title Only Author and Title

Monastyrska, I., Rieter, E., Klionsky, D.J., and Reggiori, F. (2009). Multiple roles of the cytoskeleton in autophagy. Biol Rev Camb Philos Soc 84, 431–448. Google Scholar: Author Only Title Only Author and Title

Naumann, C., Muller, J., Sakhonwasee, S., Wieghaus, A., Hause, G., Heisters, M., Burstenbinder, K., and Abel, S. (2019). The Local Phosphate Deficiency Response Activates Endoplasmic Reticulum Stress-Dependent Autophagy. Plant physiology 179, 460–476. Google Scholar: Author Only Title Only Author and Title

Nebenfuhr, A., Ritzenthaler, C., and Robinson, D.G. (2002). Brefeldin A: deciphering an enigmatic inhibitor of secretion. Plant physiology 130, 1102–1108. Google Scholar: Author Only Title Only Author and Title

Nelson, C.J., Alexova, R., Jacoby, R.P., and Millar, A.H. (2014). Proteins with high turnover rate in barley leaves estimated by proteome analysis combined with in planta isotope labeling. Plant physiology 166, 91–108. Google Scholar: Author Only Title Only Author and Title

Nesvizhskii, A.I., Keller, A., Kolker, E., and Aebersold, R. (2003). A statistical model for identifying proteins by tandem mass spectrometry. Anal Chem 75, 4646–4658. Google Scholar: Author Only Title Only Author and Title

Otegui, M.S. (2018). Vacuolar degradation of chloroplast components: autophagy and beyond. Journal of experimental botany 69, 741–750. Google Scholar: Author Only Title Only Author and Title

Pavel, M., Imarisio, S., Menzies, F.M., Jimenez-Sanchez, M., Siddiqi, F.H., Wu, X., Renna, M., O’Kane, C.J., Crowther, D.C., and Rubinsztein, D.C. (2016). CCT complex restricts neuropathogenic protein aggregation via autophagy. Nature communications 7, 13821. Google Scholar: Author Only Title Only Author and Title

Pereira, C., Pereira, S., and Pissarra, J. (2014). Delivering of proteins to the plant vacuole--an update. Int J Mol Sci 15, 7611–7623. Google Scholar: Author Only Title Only Author and Title

Qian, X., Li, X., and Lu, Z. (2017a). Protein kinase activity of the glycolytic enzyme PGK1 regulates autophagy to promote tumorigenesis. Autophagy 13, 1246–1247. Google Scholar: Author Only Title Only Author and Title

Qian, X., Li, X.J., Cai, Q.S., Zhang, C.B., Yu, Q.J., Jiang, Y.H., Lee, J.H., Hawke, D., Wang, Y.G., Xia, Y., Zheng, Y., Jiang, B.H., Liu, D.X., Jiang, T., and Lu, Z.M. (2017b). Phosphoglycerate Kinase 1 Phosphorylates Beclin1 to Induce Autophagy. Molecular cell 65, 917–+. Google Scholar: Author Only Title Only Author and Title

Razi, M., Chan, E.Y., and Tooze, S.A. (2009). Early endosomes and endosomal coatomer are required for autophagy. J Cell Biol 185, 305–321. Google Scholar: Author Only Title Only Author and Title

Reggiori, F., Monastyrska, I., Shintani, T., and Klionsky, D.J. (2005). The actin cytoskeleton is required for selective types of autophagy, but not nonspecific autophagy, in the yeast Saccharomyces cerevisiae. Mol Biol Cell 16, 5843–5856. Google Scholar: Author Only Title Only Author and Title

Rochfort, S.J., Trenerry, V.C., Imsic, M., Panozzo, J., and Jones, R. (2008). Class targeted metabolomics: ESI ion trap screening methods for glucosinolates based on MSn fragmentation. Phytochemistry 69, 1671–1679. Google Scholar: Author Only Title Only Author and Title

Schindelin, J., Arganda-Carreras, I., Frise, E., Kaynig, V., Longair, M., Pietzsch, T., Preibisch, S., Rueden, C., Saalfeld, S., Schmid, B., Tinevez, J.Y., White, D.J., Hartenstein, V., Eliceiri, K., Tomancak, P., and Cardona, A. (2012). Fiji: an open-source platform for biological-image analysis. Nat Methods 9, 676–682. Google Scholar: Author Only Title Only Author and Title

Shimada, T., Takagi, J., Ichino, T., Shirakawa, M., and Hara-Nishimura, I. (2018). Plant Vacuoles. Annu Rev Plant Biol 69, 123–145. Google Scholar: Author Only Title Only Author and Title

Shteynberg, D., Deutsch, E.W., Lam, H., Eng, J.K., Sun, Z., Tasman, N., Mendoza, L., Moritz, R.L., Aebersold, R., and Nesvizhskii, A.I. (2011). iProphet: multi-level integrative analysis of shotgun proteomic data improves peptide and protein identification rates and error estimates. Molecular & cellular proteomics : MCP 10, M111 007690. Google Scholar: Author Only Title Only Author and Title

Stobiecki, M., Skirycz, A., Kerhoas, L., Kachlicki, P., Muth, D., Einhorn, J., and Mueller-Roeber, B. (2006). Profiling of phenolic glycosidic conjugates in leaves of Arabidopsis thaliana using LC/MS. Metabolomics 2, 197–219. Google Scholar: Author Only Title Only Author and Title

Styers, M.L., O’Connor, A.K., Grabski, R., Cormet-Boyaka, E., and Sztul, E. (2008). Depletion of beta-COP reveals a role for COP-I in compartmentalization of secretory compartments and in biosynthetic transport of caveolin-1. Am J Physiol Cell Physiol 294, C1485–1498. Google Scholar: Author Only Title Only Author and Title

Sugiyama, R., and Hirai, M.Y. (2019). Atypical Myrosinase as a Mediator of Glucosinolate Functions in Plants. Frontiers in plant science 10. Google Scholar: Author Only Title Only Author and Title

Tasaki, M., Asatsuma, S., and Matsuoka, K. (2014). Monitoring protein turnover during phosphate starvation-dependent autophagic degradation using a photoconvertible fluorescent protein aggregate in tobacco BY-2 cells. Frontiers in plant science 5, 172. Google Scholar: Author Only Title Only Author and Title

Thompson, A.R., Doelling, J.H., Suttangkakul, A., and Vierstra, R.D. (2005). Autophagic nutrient recycling in Arabidopsis directed by the ATG8 and ATG12 conjugation pathways. Plant physiology 138, 2097–2110. Google Scholar: Author Only Title Only Author and Title

Thormahlen, I., Zupok, A., Rescher, J., Leger, J., Weissenberger, S., Groysman, J., Orwat, A., Chatel-Innocenti, G., Issakidis-Bourguet, E., Armbruster, U., and Geigenberger, P. (2017). Thioredoxins Play a Crucial Role in Dynamic Acclimation of Photosynthesis in Fluctuating Light. Mol Plant 10, 168–182. Google Scholar: Author Only Title Only Author and Title

Vandesompele, J., De Preter, K., Pattyn, F., Poppe, B., Van Roy, N., De Paepe, A., and Speleman, F. (2002). Accurate normalization of real-time quantitative RT-PCR data by geometric averaging of multiple internal control genes. Genome Biol 3, RESEARCH0034. Google Scholar: Author Only Title Only Author and Title

Vincow, E.S., Thomas, R.E., Merrihew, G.E., Shulman, N.J., Bammler, T.K., MacDonald, J.W., MacCoss, M.J., and Pallanck, L.J. (2019). Autophagy accounts for approximately one-third of mitochondrial protein turnover and is protein selective. Autophagy 15, 1592–1605. Google Scholar: Author Only Title Only Author and Title

Wang, J., Zhang, J., Lee, Y.M., Koh, P.L., Ng, S., Bao, F., Lin, Q., and Shen, H.M. (2016). Quantitative chemical proteomics profiling of de novo protein synthesis during starvation-mediated autophagy. Autophagy 12, 1931–1944. Google Scholar: Author Only Title Only Author and Title

Wang, S., Xie, K., Xu, G., Zhou, H., Guo, Q., Wu, J., Liao, Z., Liu, N., Wang, Y., and Liu, Y. (2018). Plant G proteins interact with endoplasmic reticulum luminal protein receptors to regulate endoplasmic reticulum retrieval. J Integr Plant Biol 60, 541–561. Google Scholar: Author Only Title Only Author and Title

Waters, M.T., Nelson, D.C., Scaffidi, A., Flematti, G.R., Sun, Y.K., Dixon, K.W., and Smith, S.M. (2012). Specialisation within the DWARF14 protein family confers distinct responses to karrikins and strigolactones in Arabidopsis. Development 139, 1285–1295. Google Scholar: Author Only Title Only Author and Title

Watson, A.S., Riffelmacher, T., Stranks, A., Williams, O., De Boer, J., Cain, K., MacFarlane, M., McGouran, J., Kessler, B., Khandwala, S., Chowdhury, O., Puleston, D., Phadwal, K., Mortensen, M., Ferguson, D., Soilleux, E., Woll, P., Jacobsen, S.E., and Simon, A.K. (2015). Autophagy limits proliferation and glycolytic metabolism in acute myeloid leukemia. Cell Death Discov 1. Google Scholar: Author Only Title Only Author and Title

Wijerathna-Yapa, A., Stroeher, E., Fenske, R., Li, L., Duncan, O., and Millar, A.H. (2021). Proteomics for Autophagy Receptor and Cargo Identification in Plants. J Proteome Res 20, 129–138. Google Scholar: Author Only Title Only Author and Title

Xiong, Y., Contento, A.L., and Bassham, D.C. (2005). AtATG18a is required for the formation of autophagosomes during nutrient stress and senescence in Arabidopsis thaliana. Plant J 42, 535–546. Google Scholar: Author Only Title Only Author and Title

Xu, J., Kozlov, G., McPherson, P.S., and Gehring, K. (2018). A PH-like domain of the Rab12 guanine nucleotide exchange factor DENND3 binds actin and is required for autophagy. J Biol Chem 293, 4566–4574. Google Scholar: Author Only Title Only Author and Title

Yokota, H., Gomi, K., and Shintani, T. (2017). Induction of autophagy by phosphate starvation in an Atg11-dependent manner in Saccharomyces cerevisiae. Biochem Biophys Res Commun 483, 522–527. Google Scholar: Author Only Title Only Author and Title

Yoshimoto, K., Ishida, H., Wada, S., Ohsumi, Y., and Shirasu, K. (2009a). The role of plant autophagy in nutrient starvation and aging. Autophagy 5, 904–904. Google Scholar: Author Only Title Only Author and Title

Yoshimoto, K., Jikumaru, Y., Kamiya, Y., Kusano, M., Consonni, C., Panstruga, R., Ohsumi, Y., and Shirasu, K. (2009b). Autophagy negatively regulates cell death by controlling NPR1-dependent salicylic acid signaling during senescence and the innate immune response in Arabidopsis. The Plant cell 21, 2914–2927. Google Scholar: Author Only Title Only Author and Title

Zhang, C., Hicks, G.R., and Raikhel, N.V. (2014). Plant vacuole morphology and vacuolar trafficking. Frontiers in plant science 5. Google Scholar: Author Only Title Only Author and Title

Zhang, K., Halitschke, R., Yin, C., Liu, C.J., and Gan, S.S. (2013). Salicylic acid 3-hydroxylase regulates Arabidopsis leaf longevity by mediating salicylic acid catabolism. Proc Natl Acad Sci U S A 110, 14807–14812. Google Scholar: Author Only Title Only Author and Title

Zhang, T., Shen, S., Qu, J., and Ghaemmaghami, S. (2016). Global Analysis of Cellular Protein Flux Quantifies the Selectivity of Basal Autophagy. Cell Reports 14, 2426–2439. Google Scholar: Author Only Title Only Author and Title

Zhang, X., Ding, X., Marshall, R.S., Paez-Valencia, J., Lacey, P., Vierstra, R.D., and Otegui, M.S. (2020). Reticulon proteins modulate autophagy of the endoplasmic reticulum in maize endosperm. Elife 9. Google Scholar: Author Only Title Only Author and Title

Zheng, X., Wu, M., Li, X., Cao, J., Li, J., Wang, J., Huang, S., Liu, Y., and Wang, Y. (2019). Actin filaments are dispensable for bulk autophagy in plants. Autophagy 15, 2126–2141. Google Scholar: Author Only Title Only Author and Title

Zhuang, X., and Jiang, L. (2019). Chloroplast Degradation: Multiple Routes Into the Vacuole. Frontiers in plant science 10, 359. Google Scholar: Author Only Title Only Author and Title

